# Interplay Between FoxM1 and Dab2 Promotes Endothelial Cell Responses in Diabetic Wound Healing

**DOI:** 10.1101/2024.02.07.579019

**Authors:** Sudarshan Bhattacharjee, Jianing Gao, Yao Wei Lu, Shahram Eisa-Beygi, Hao Wu, Kathryn Li, Amy E. Birsner, Scott Wong, Yudong Song, John Y-J. Shyy, Douglas B. Cowan, Wenyi Wei, Masanori Aikawa, Jinjun Shi, Hong Chen

## Abstract

Diabetes mellitus can cause impaired and delayed wound healing, leading to lower extremity amputations; however, the mechanisms underlying the regulation of vascular endothelial growth factor (VEGF)-dependent angiogenesis remain uncertain and could reveal new therapeutic targets. In our study, the molecular underpinnings of endothelial dysfunction in diabetes were investigated, focusing on the roles of Disabled-2 (Dab2) and Forkhead Box M1 (FoxM1) in VEGF receptor 2 (VEGFR2) signaling and endothelial cell (EC) function. Bulk RNA-sequencing analysis identified significant downregulation of Dab2 in high concentrations glucose treated primary mouse skin ECs, simulating hyperglycemic conditions in diabetes mellitus. In diabetic mice with a genetic EC deficiency of Dab2 angiogenesis was reduced *in vivo* and *in vitro* when compared with wild-type mice. Restoration of Dab2 expression by injected mRNA-containing lipid nanoparticles rescued impaired angiogenesis and wound healing in diabetic mice. At the same time, FoxM1 was downregulated in skin ECs subjected to high glucose conditions as determined by RNA-sequencing analysis. FoxM1 was found to bind to the Dab2 promoter, regulating its expression and influencing VEGFR2 signaling. The FoxM1 inhibitor FDI-6 reduced Dab2 expression and phosphorylation of VEGFR2. These findings indicate that restoring Dab2 expression through targeted therapies can enhance angiogenesis and wound repair in diabetes. To explore this therapeutic potential, we tested LyP-1-conjugated lipid nanoparticles (LNPs) containing Dab2 or control mRNAs to target ECs and found the former significantly improved wound healing and angiogenesis in diabetic mice. This study provides evidence of the crucial roles of Dab2 and FoxM1 in diabetic endothelial dysfunction and establishes targeted delivery as a promising treatment for diabetic vascular complications.

## Introduction

One of the most serious pathological outcomes of diabetes mellitus is impaired and/or delayed wound healing, which – in severe cases – can lead to lower extremity amputations (LEAs)^1–3^. Although the etiological basis of chronic non-healing wounds is multi-faceted, aberrant angiogenesis is, at least in part, involved in sustaining this phenotype. During wound healing, angiogenic sprouts descend upon the wound area to establish normoxia and they eventually fashion a microvascular network to restore oxygen and nutrient delivery to the wound area and help remove debris^4–6^. Therefore, promoting angiogenesis is crucial for wound healing, and developing effective targets for angiogenesis could benefit millions of diabetic patients.

Vascular endothelial growth factor (VEGF) is a critical angiogenic factor that signals through VEGF receptors (VEGFRs)^7^. Among the family of VEGFRs, VEGFR2 potentiates angiogenesis more potently than other VEGFRs. Binding of VEGF to VEGFR2 leads to the phosphorylation of VEGFR2 and activation of downstream signaling pathways, including MAPK/ERK and PI3K/Akt, which promote endothelial cell (ECs) proliferation, migration, and survival^8,9^. In diabetic conditions, there is a decrease in VEGF-induced phosphorylation of VEGFR2 and downstream signaling, leading to impaired angiogenesis^10–12^. Hence, gaining insights into the regulation of VEGFR2-dependent angiogenesis may lead to the identification of new therapeutic strategies in this context.

Several studies have shown that the highly-conserved adaptor protein Disabled Homolog 2 (Dab2) plays a direct role in regulating VEGF signaling in ECs^13,14^. Dab2 is involved in the regulation of endocytosis and lysosomal degradation of receptor tyrosine kinases, including VEGFR2. Dab2 binds to the cytoplasmic tail of VEGFR2 and promotes its endocytosis and recycling^14^; thereby, serving to enhance VEGFR2-mediated angiogenesis. In spite of this, the specific structural domain through which Dab2 interacts with VEGFR2 is unknown. At the same time, the molecular mechanisms underlying Dab2-mediated angiogenesis, particularly in the context of wound healing in diabetes, is not clear. Identifying factors that regulate Dab2 transcription is key to uncovering the mechanisms of Dab2’s role in endothelial cell angiogenesis under diabetic conditions. More pressingly, it remains unknown if modifying Dab2 levels could serve as a therapeutic strategy to enhance diabetic wound healing. To better target Dab2, it is essential to develop exogenous supplementation methods that have a shorter half-life and higher efficiency.

The present study was designed to dissect the potential involvement of Dab2 in regulating VEGF signaling during angiogenesis in the context of wound healing in diabetes. Using an EC-specific Dab2 knockout mouse model, we found that the Forkhead box M (FoxM1) regulates Dab2 expression in ECs and promotes diabetic wound healing. This transcription factor orchestrates the expression of genes essential for cell cycle progression, thus facilitating cell growth and division; a process that is vital for tissue repair and regeneration^15–18^. We found that FoxM1 positively regulates Dab2 expression by directly binding to its promoter to influence transcription and protein levels of Dab2. By injecting Dab2-mRNA encapsulated in lipid nanoparticles (LNPs), or using the FoxM1 inhibitor FDI-6, wound healing was significantly enhanced through increased angiogenesis, which could lay the foundation for the development of novel therapies to enhance angiogenesis in diabetes.

Together, our results suggest that Dab2 plays a critical role in VEGF signaling and angiogenesis in ECs under diabetic conditions by regulating the activation of VEGFR2. We also identified the specific binding domain of Dab2 that binds to VEGFR2 and demonstrated that FoxM1 regulated transcription of this adaptor protein in ECs. Our findings indicate that Dab2 may represent a previously unidentified potential target for improving diabetic wound healing.

## Methods

### Mouse models

All animal experiments were approved by the Institutional Animal Care and Use Committee at Boston Children’s Hospital. To produce EC-specific Dab2KO (Dab2-EC^iKO^) mice, a breeding strategy was employed using Dab2^fl/fl^ mice and EC-specific Cre transgenic CDH5-Cre mice. To activate Cre recombinase, 8 to 10 week old mice received 4-hydroxytamoxifen (Hello Bio, dissolved in a 9:1 mixture of DMSO and ethanol at a dosage of 5–10 mg/kg body weight) seven times every other day. For induction of diabetes, mice underwent intraperitoneal injection with a low-dose of streptozotocin (STZ, Sigma-Aldrich, 50 mg/kg) following an established protocol^19^. Hyperglycemia was confirmed when mice maintained a fasting blood glucose level above 200 mg/dL for over a week post-STZ administration. After inducing diabetes, the mice were placed on a high-fat diet (HFD, 60 kcal% fat from Research Diets Inc.).

### Cell cultures

Primary mouse ECs were obtained from mouse skin and cultured according to previously established protocols with some modifications^19–21^. Briefly, to isolate ECs from the skin, 4-6 mice aged 2-3 months were used. The mice were anesthetized with isoflurane and humanely euthanized by cervical dislocation. The skin was excised from the mice using surgical scissors or a scalpel, then diced into small fragments on ice and subjected to enzymatic digestion with collagenase Type IV (2 mg/mL; Gibco Laboratories) in a 37°C water bath with agitation (10 rpm) for a duration of 60-90 minutes. Digestion was stopped by adding an equal volume of ice-cold FBS. The resulting digested tissue was filtered through a 40 µm cell strainer (BD) to separate the cells from debris. The resulting cell suspension was then centrifuged at 400 g for 5 minutes at 4°C. 10 μL anti-mouse CD31 MicroBeads (Miltenyi Biotec) were added into about 10^7^ isolated cells in 90 μL of buffer (PBS, pH 7.2, 0.5% BSA, and 2 mM EDTA). The cell mixture was then incubated for 15 minutes at 4 °C. These isolated cells were utilized for downstream experiments. The primary ECs used in all experiments were isolated and maintained between 1-6 passages. Cells were treated with normal glucose (5 mmol/L) or high glucose (20 mmol/L) medium for ∼48 hours. ECs derived from wild-type mice or mice carrying Dab2^fl/fl^; iCDH5-ER^T2^ Cre alleles were exposed to 5 μmol/L of 4-hydroxytamoxifen (dissolved in ethanol) for two days at 37°C. Following treatment, cells were incubated for another two days without 4-hydroxytamoxifen. Confirmation of Dab2 deletion was carried out by Western Blotting.

### Mouse corneal micropocket angiogenesis assay

The corneal micropocket angiogenesis assay in mice was carried out following established protocols^19,22^. Briefly, mice were anesthetized using Avertin (400-500 mg/kg delivered by intraperitoneal injection i.p.). An incision into the cornea was gently created at an approximately 30° angle and 0.7 -1.0 mm from the limbus using a corneal blade and a stereoscope. A sustained-release pellet containing the volume of 0.4 mm x 0.4 mm x 0.2 mm pellet of VEGFA (∼ 20 ng, BioLegend) was implanted into the pocket. 5 -7 days post-implantation, the corneas were excised, and stained with PE-conjugated anti-CD31 antibody (1:100, BD Pharmingen) to highlight limbal blood vessels. The growth of these vessels was quantified by measuring the growth pixels using the Vessel Analysis plugin in ImageJ.

### RNA isolation

Total RNA was isolated from the cells using a commercially-available kit (Qiagen, Valencia, CA, USA) according to the manufacturer’s instructions. RNA quantity and quality was determined using a NanoDrop 2000 spectrophotometer (Thermo Fisher Scientific, Waltham, MA, USA).

### Library preparation and bulk RNA sequencing

Total RNA was extracted from the collected samples and assessed for quality and integrity. Quality control of all RNA-seq samples was performed at the Harvard Medical School Biopolymer Facility using a 2100 Bioanalyzer (Agilent). Samples with an RIN score greater than 7 were considered for further processing. The NEBNext Poly(A) mRNA Magnetic Isolation Module was used to enrich poly(A)+ mRNA from the total RNA pool, using Oligo d(T)25 beads for magnetic separation. Post isolation, the mRNA was eluted and either used immediately or stored at -80°C for subsequent experiments. Library preparation began with reverse transcription of the enriched mRNA into cDNA, followed by end-repair and the addition of a single ‘A’ nucleotide at the 3’ ends. NEBNext Adaptors were then ligated to the cDNA, and unique indices were added using the NEBNext Multiplex Oligos for Illumina to allow for sample multiplexing during sequencing. The adaptor-ligated cDNA was used for PCR amplification, and the product was purified to remove any remaining enzymes and primers. The final library was validated for quality using capillary electrophoresis and quantified through qPCR. Libraries were pooled in equimolar ratios, as determined by the quantification results, before being forwarded to the sequencing facility for high-throughput analysis. The RNA library was sequenced using the HiSEQ Next-generation Sequencing System with a read length of 75 at the core facility of Azenta Life Sciences.

### Bulk RNA-seq data analysis

Sequencing data were processed using Trimmomatic software. Low-quality reads were removed to retain only high-quality sequences for analysis. These clean reads were then aligned to the mouse reference genome GRCm38 (mm10) using the HISAT2 alignment tool. StringTie was used to enumerate the reads for each gene. The edgeR package in R was used to identify genes with significant changes in expression, with significant differences being noted at an adjusted p-value of less than 0.05. Pathway and gene function analyses were performed using Gene Set Enrichment Analysis (GSEA), with significance attributed to terms with an adjusted p-value and FDR below 0.05. Genes were categorized in the volcano plot based on the degree of expression change and statistical significance, with colors assigned accordingly.

### qRT-PCR

Total RNA was isolated from primary mouse ECs using a commercially available kit (Qiagen, Valencia, CA, USA) followed by DNase I treatment. The RNA was then reverse transcribed into cDNA using Oligo (dT) 20 Primers (Invitrogen) as per the manufacturer’s protocol. Quantitative real-time PCR was performed using the StepOnePlus Real-Time PCR detection system (Applied Biosystems, Foster City, CA, USA) and SYBR Green qPCR super Mix (Invitrogen). The PCR amplification cycles consisted of an initial heating step at 95°C for 10 min, followed by 40 cycles of 15 s at 95°C, 1 minute at 60°C, and 45 s at 72°C. The relative abundance of mRNA was determined using the average threshold cycles (Ct) of samples normalized to β-actin mRNA. Each sample was analyzed in triplicate. The mouse primer sequences used for PCR are shown in Table S1.

### RNA Interference

The study used RNA interference (RNAi) to knockdown the expression of specific genes in primary ECs. ON-TARGETplus Mouse Dab2 siRNA (Horizon, J-050859-09-0002) and its respective ON-TARGETplus non-targeting siRNAs (Horizon, D-001810-0X) were transfected in the isolated ECs using either Oligofectamine or RNAiMAX according to the manufacturer’s instructions (Invitrogen). The cells were processed for biochemical immunoprecipitations or immunofluorescence assays 48-72 hours after transfections as previously described ^23^.

### Glucose tolerance test (GTT)

Glucose tolerance was tested as described previously^19^. Mice were fasted for 16 hours. The weight of each mouse determined the calculated glucose dose at a ratio of 2 g/kg body weight. Injection volume into the peritoneal of mouse = BW(g)×10μL of 250mg/mL glucose solution. Tail snipping was used to collect blood samples. Blood glucose was determined by glucometer in tail vein blood. Blood glucose is measured at 0, 30, 60, 90, and 120 minutes after glucose injection.

### Insulin tolerance test (ITT)

Insulin tolerance was tested as described previously^19^. Mice are fasted for 3 hours. The weight of each mouse determined the calculated insulin dose at a ratio of 0.75 U/kg body weight. Insulin was prepared at 0.1 U/mL in advance (16.6μL of 10 mg/mL insulin in 40 mL PBS). Injection volume into the peritoneal of mouse = BW(g) X 7.5μL of 0.1U/mL insulin solution. Tail snipping is used to obtain blood and glucose levels were determined using a glucometer. Measurements were made at 0, 15, 30, 60, 90, and 120 minutes after insulin injection.

### LyP-1 peptide-linked Dab2 and control mRNA lipid nanoparticles

LyP-1 peptide was conjugated to DSPE-PEG via an NHS-amine reaction^24–27^. Briefly, the activated DSPE-PEG-NHS was mixed with Lyp1 in 1x PBS buffer solution (pH 7.4) at room temperature and stirred for 24 hours. These crude products were dialyzed against water for 3 days (MWCO, 3 kDa), followed by lyophilization. Successful conjugation was confirmed using proton nuclear magnetic resonance (1H NMR)^28^.

Lipid nanoparticles (LNPs) containing mRNA, including LNPs-Lyp1-Dab2 mRNA and LNPs-Lyp1-GFP mRNA were formulated by mixing an aqueous phase containing mRNA and an organic ethanol phase containing MC3, 1,2-dioleoyl-sn-glycero-3-phosphoethanolamine (DOPE), cholesterol, and PEG-conjugated lipids (DMG-PEG and DSPE-PEG) ^29,30^. Briefly, one volume of lipid mixtures (MC3, DOPE, Chol, DMG-PEG, and DSPE-PEG-LyP1 at a molar ratio of 50:10:38: 1:1) in ethanol and three volumes of mRNA (Dab2 mRNA or GFP mRNA, 1:10 w/w mRNA to lipid) containing sodium acetate buffer (50 mM, pH 4) were mixed thoroughly and stirred at RT for 20 min. The resulting LNPs-Lyp1-Dab2 mRNA and LNPs-Lyp1-GFP mRNA were further purified by ultrafiltration (MWCO, 10 kDa) with 1x PBS (pH7.4) to remove naked mRNAs and ethanol. The final mRNA-loaded LNPs were maintained in 1x PBS at an mRNA concentration of 75 µg/mL^30^. Dynamic light scattering (DLS) was adopted to characterize the LNPs-Lyp1-Dab2 mRNA and LNPs-Lyp1-GFP mRNA^29,31^. As shown in Fig. S3B, the average sizes were 136.7 ± 2.52 nm for LNPs-Lyp1-Dab2 mRNA and 120.5 ± 2.45 nm for LNPs-Lyp1-GFP mRNA, with zeta potentials of -5.54 ± 0.17 mV and -6.11 ± 0.51 mV, respectively.

### Injection of LNPs into mice

Dab2 mRNA was loaded into LNPs to facilitate the *in vivo* application by protecting mRNA from enzymatic degradation, enhancing cellular uptake and endosomal escape, and/or improving systemic circulation. This encapsulation process was crucial to maintain the stability and effectiveness of the Dab2 mRNA for its use in living organisms. The prepared Dab2-mRNA LNPs (LNPs-Lyp1-Dab2 mRNA or control LNPs-Lyp1-GFP mRNA) were administered to mice by intravenous (i.v.) injection, with a dosage of 15 μg per mouse. These treatments were given twice a week for a total of four weeks.

### Lentivirus-mediated Dab2 overexpression

Lentivirus for Dab2 overexpression was produced using the PEI STAR transfection method ^32–35^ in ECs. Initially, ECs were seeded in a 10 cm dish and incubated at 37°C with 5% CO_2_ until they reached 60-70% confluence. For transfection, a mixture containing equimolar amounts of pMD2.G (VSVg) and pSPAX2 (Gag and Pol) and, along with a double molar amount of pGenLenti Dab2-Flag transfer vector was prepared for a total of 10 µg of DNA. This DNA mixture was combined with 30 µL of PEI STAR (1 mg/mL) in 500 µL of fresh medium, and incubated for 10 minutes, and then added dropwise to the cells. After 24 hours, the medium was replaced to remove residual transfection reagent, and the process was repeated after 48 hours. After 72 hours of transfection, the supernatant containing the viral particles was harvested and centrifuged to remove cell debris, which was filtered through a 0.45 µm filter and stored at -80°C for subsequent experiments.

### CRISPR/Cas9-mediated mutations to block FoxM1 binding to the mouse *Dab2* promoter

CRISPR/Cas9-mediated gene editing was employed to introduce targeted mutations within the Dab2 promoter region of mouse skin ECs to disrupt the FoxM1 binding site. The Ad5CMVspCas9/RSVeGFP vector was purchased from the Viral Core at the University of Iowa, using the expression of SpCas9 and a GFP reporter. The selected sgRNA sequences: sgRNA1: TAAGATTCTCTACTATGTG (+ Strand); sgRNA2: TTGTATATATCTTGGGGAA (-Strand). A 1,154 bp DNA fragment with three FoxM1 transcription factor binding sites spans from -4,806 bp to -5,906 bp upstream of the transcription start site (TSS) of Dab 2. The synthetic DNA fragment containing mutations in the PAMs and FoxM1 binding sites was obtained from Synthego. To prevent FoxM1 from binding to the Dab2 promoter and verify whether cell transduction is successful, the three binding sites of FoxM1 were mutated into restriction enzyme sites in the synthetic DNA fragment, as follows: Site 1: Sal1 restriction site (AAATGC -> GTCGAC); site 2: Fsp1 restriction site (CAATGC -> TGCGCA); site 3: Sal1 restriction site (TAATGA -> GTCGAC).

For the transduction of ECs, the Ad5CMVspCas9/RSVeGFP vector was introduced using Lipofectamine 3000 (Thermo Fisher Scientific) to maintain the integrity and viability of cells. After 3 days, the synthetic DNA recombination fragment DNA fragment was introduced into the ECs employing the Amaxa Nucleofector 1 Electroporation System and associated kit, strictly adhering to the manufacturer’s protocol. The electroporation conditions were optimized to ensure high viability and efficient uptake of the DNA constructs. After transfection, cells were cultured under standard conditions and screened for GFP expression using fluorescence microscopy. DNA isolated from transduced EC cells was then digested with restriction enzymes to further confirm successful delivery and expression of CRISPR components.

### ChIP-PCR

Following genomic editing, ChIP-PCR was performed to verify the impact of mutation on FoxM1 binding to the Dab2 promoter. The ChIP procedure made use of a kit (Abcam), starting with cell fixation, chromatin shearing by sonication, and immunoprecipitation with specific antibodies targeting FoxM1. The DNA-protein complexes were pulled down using protein A beads, and the DNA was purified and analyzed by PCR to assess the binding activity of FoxM1 at the modified Dab2 promoter. The ChIP-PCR primers of Dab2 are as follows: Forward: 5’-CCCAGCAGTACAAGTCTGGA-3’; Reverse: 5’-AGGACTGAGTGGACATGGTG-3’.

### Western blot analysis

To extract total proteins, cells were lysed using RIPA buffer. Equal amounts of denatured protein were loaded onto a 10% SDS-PAGE gel for electrophoresis. The separated proteins were then transferred to a PVDF membrane and blocked with 5% skimmed milk for 30 minutes at room temperature. Primary antibodies were diluted at 1:1000 and the secondary antibodies were diluted at 1:2000. The antibodies are listed in Table S2.

### Immunofluorescence staining

For immunofluorescence staining, primary mouse ECs were cultured on glass coverslips and fixed in 4% paraformaldehyde (PFA) for 10 minutes. The cells were then permeabilized with 0.3% Triton X-100 in phosphate-buffered saline (PBS) for 10 minutes and blocked in a solution containing 5% donkey serum in PBS for 1 hour at room temperature. The primary antibodies were diluted in blocking solution and incubated with the cells overnight at 4°C. The next day, the coverslips were washed with PBS and incubated with the respective secondary antibodies conjugated to fluorescent labels (Alexa Fluor) for 2 hours at room temperature. After washing with PBS, the coverslips were mounted on glass slides using Vectashield mounting medium with DAPI and visualized using a fluorescence microscope (Zeiss LSM 880) with appropriate filters. Images were acquired using a digital camera (Zeiss AxioCam). Unless otherwise specified, secondary antibodies for immunohistochemistry were applied at a concentration of 1:200. The antibodies are listed in Table S2.

### *In vitro* wound healing assays

An *in vitro* scratch wound healing assay was performed to evaluate the migration of wild type and Dab2-EC^iKO^ cells under normoglycemic and hyperglycemic conditions. Cells were seeded in a 24-well plate and grown to confluence. A scratch wound was made in the cell monolayer using a sterile pipette tip, and the cells were washed to remove any detached cells. The cells were then cultured in either a normoglycemic media (5 mmol/L glucose) or a hyperglycemic media (20 mmol/L glucose) in the presence or absence of VEGFA (100ng/mL) and images were taken at specified time points to monitor cell migration into the wound area. The rate of cell migration was calculated by measuring the width of the wound at different time points. The differences in wound closure between wild type and Dab2-EC^iKO^ cells under both normoglycemic and hyperglycemic conditions were analyzed.

### Vascular network formation assays

An *in vitro* Matrigel assay was performed to compare the behavior of wild type and Dab2-EC^iKO^ cells in normoglycemic and hyperglycemic conditions. The cells were seeded on top of a Matrigel matrix in a 24-well plate and cultured in a normoglycemic media (5 mmol/L glucose) or a hyperglycemic media (20 mmol/L glucose) for a specified period. The cells were fixed and analyzed to determine morphological changes and quantify cell migration and proliferation. The differences in behavior between the wild type and Dab2-EC^iKO^ cells were analyzed under both normoglycemic and hyperglycemic conditions.

### EdU staining

An EdU staining assay was performed to determine the cell proliferation of wild type and Dab2-EC^iKO^ cells under normoglycemic and hyperglycemic conditions. The cells were cultured in either a normoglycemic media (5 mmol/L glucose) or a hyperglycemic media (20 mmol/L glucose) and then incubated with EdU, a thymidine analog, for 4 hours in presence or absence of VEGFA (100 ng/mL). The cells were then fixed and processed for staining, including the addition of a DAPI stain to visualize the nuclei. The EdU-incorporated cells were identified by fluorescence using a specific antibody. The number of EdU positive cells was counted and compared between the wild type and Dab2-EC^iKO^ cells under both normoglycemic and hyperglycemic conditions. The differences in cell proliferation were analyzed and quantified.

### *In vivo* wound healing assays

A dermal wound healing assay was performed to evaluate the role of the Dab2 gene in the wound healing process. Two groups of mice, wild type and Dab2-EC^iKO^, were used in this study. A standardized wound was created on the dorsal surface of each mouse using a surgical blade. The mice were then monitored for approximately 7 days to evaluate wound healing, including changes in wound size and tissue regeneration. At the end of the study, the mice were sacrificed, and the wound sites were collected for further analysis. The collected tissue was processed for histological analysis, including the use of specific stains to evaluate vascular density during wound closure. The differences in wound healing response between the wild type, diabetic, and Dab2-EC^iKO^ mice were analyzed and quantified.

### Matrigel plug assays

An *in vivo* Matrigel assay was performed to study the angiogenic response of wild type and Dab2-EC^iKO^ mice. Matrigel mixed with 100 ng/mL VEGFA was prepared, and 400 - 500 µL was implanted subcutaneously into the back of both wild type and Dab2-EC^iKO^ mice. The mice were then monitored for 7 days to evaluate angiogenic response, including the formation of blood vessels in the implantation site. At the end of the study, the mice were sacrificed, and the implantation sites were collected for further analysis. The collected tissue was processed for histological analysis, including the use of specific stains to visualize blood vessels and quantify angiogenic response. The differences in angiogenic response between the wild type and Dab2-EC^iKO^ mice were analyzed and quantified. This assay provided valuable information on the role of the Dab2 gene in modulating angiogenic response *in vivo*.

### Dab2 inhibitors predicted by molecular modeling

To predict the Dab2 signaling inhibitors, docking experiments were performed using the ClusPro 2.0 program^36,37^. The 3-D structure of Dab2 PTB domain (PDB ID: 2LSW) was docked into VEGFR2 Kinase domain (PDB ID: 3U6J) to generate the predicted binding models of Dab2:VEGFR2 (Table 1). Models with the highest scores and best topologies were selected for the proposed models of the interaction between Dab 2 and VEGFR2. In the interaction models, a total of 6 Dab2 inhibitory peptides were identified with good scores based on the molecular modeling (Fig. 3H-I).

**Table 1.** Interacting residues between mouse Dab2 (2LSW) and VEGFR2. The crystal structure of mouse Dab2 (PDB ID: IP3R) and VEGFR2 were taken from the PDB database and used to perform the docking experiments using ClusPro 2.0. There are total 200 models per each docking experiment. The frequency of residues in Dab2 and VEGFR2 to form H-bonds among these models were ranked, respectively. From the modeling, the E33, K49, Q153, E37, and Q88 residues from Dab2 and the R1027, K1023, R1126, D1129, R819, R1022, H816, and Y996 residues from VEGFR2 were critical to form the complex.

| Interaction residues of Fibromodulin to form H-bonds |  | Interaction residues of AXL to form H-bonds |  |
| --- | --- | --- | --- |
| Predicted residues in Dab2(IP3R) | Frequency to form H-bonds | Predicted residues in VEGFR2 | Frequency to form H-bonds |
| E33 | 21 | R1027 | 27 |
| K49 | 17 | K1023 | 15 |
| Q153 | 17 | R1126 | 15 |
| W37 | 16 | D1129 | 14 |
| M32 | 16 | R819 | 13 |
| Q88 | 12 | R1022 | 11 |
| E156 | 11 | H816 | 11 |
| K108 | 11 | Y996 | 11 |
| R42 | 10 | R1172 | 10 |
| N131 | 10 | E815 | 9 |
| Q154 | 10 | K826 | 9 |
| Y38 | 9 | I170 | 9 |
| R92 | 9 | R1080 | 9 |
| Q86 | 9 | D1046 | 8 |
| R132 | 9 | N933 | 8 |
| K44 | 8 | H1173 | 7 |
| T35 | 8 | H891 | 6 |
| H89 | 7 | R932 | 6 |
| R84 | 7 | Q1149 | 6 |
| E113 | 7 | E1146 | 5 |
| K108 | 7 | E1134 | 5 |
| I57 | 7 | D1171 | 5 |
| R126 | 6 | Y1082 | 5 |
| D106 | 5 | Q1137 | 4 |
| Q91 | 5 | 935 | 4 |
| D164 | 5 | L995 | 4 |
| T109 | 5 | Y1130 | 4 |
| D59 | 4 | R1061 | 3 |
| D161 | 4 | D1028 | 3 |
| Y50 | 4 | E993 | 3 |
| K34 | 4 | R1124 | 3 |
| E115 | 4 | D1141 | 3 |
| Q145 | 4 | D994 | 3 |
| Q143 | 4 | D823 | 3 |
| K163 | 3 | R880 | 3 |
| H114 | 3 | I1025 | 3 |
| D66 | 3 | G1122 | 3 |
| K90 | 3 | H1159 | 2 |
| D46 | 3 | E1155 | 2 |
| K51 | 3 | Q1085 | 2 |
| T129 | 3 | E1158 | 2 |
| G87 | 3 | Y1054 | 2 |
| S85 | 3 | R1066 | 2 |
| K53 | 2 | H894 | 2 |
| K75 | 2 | Y938 | 2 |
| K77 | 2 | Y1136 | 2 |
| K150 | 2 | R1118 | 2 |
| R64 | 2 | R1022 | 2 |
| Q91 | 2 | Y822 | 1 |
| Q167 | 2 | H1026 | 1 |
|  |  | D1064 | 1 |
|  |  | R1051 | 1 |
|  |  | R1052 | 1 |
|  |  | K1120 | 1 |
|  |  | E815 | 1 |
|  |  | E1017 | 1 |
|  |  | R1051 | 1 |
|  |  | R1052 | 1 |
|  |  | K997 | 1 |
|  |  | K1070 | 1 |

### Statistics

Results were presented as the mean ± SD. The evaluation of differences between groups was performed using Student’s t-test with the demonstration of homogeneity of variance. The one-way ANOVA was used for multiple comparisons followed by Dunnett’s post hoc analysis, by GraphPad Prism 8. A p-value of less than 0.05 was considered for statistical significance.

## Results

### Diabetes and high glucose treatment in ECs leads to the downregulation of Dab2

Given the crucial roles in both physiological and pathological angiogenesis, we sought to determine if endocytic adaptor proteins were also involved in mitigating aspects of diabetic angiogenesis. To address this goal, we isolated CD31-enriched primary mouse endothelial cells (ECs) and treated them with normal (5 mmol/L) or high glucose concentrations (20 mmol/L), a condition mimicking hyperglycemia in diabetes mellitus, for a period of 48 hours, followed by bulk RNA-sequencing analysis. Differential gene expression analysis revealed 168 significantly downregulated and 386 significantly up-regulated genes in the ECs grown in high glucose culture conditions compared to ECs cultured in normal glucose media (Fig. 1A, S1A). Volcano plot analysis revealed downregulation of the Dab2 mRNA levels in high glucose-treated ECs compared to the control group (Fig. 1A). Further corroborating these findings, quantitative PCR (qPCR) analysis showed that high glucose treatment or ECs from diabetic mice resulted in the downregulation of Dab2 (Fig. 1B-C).

**Figure 1.**
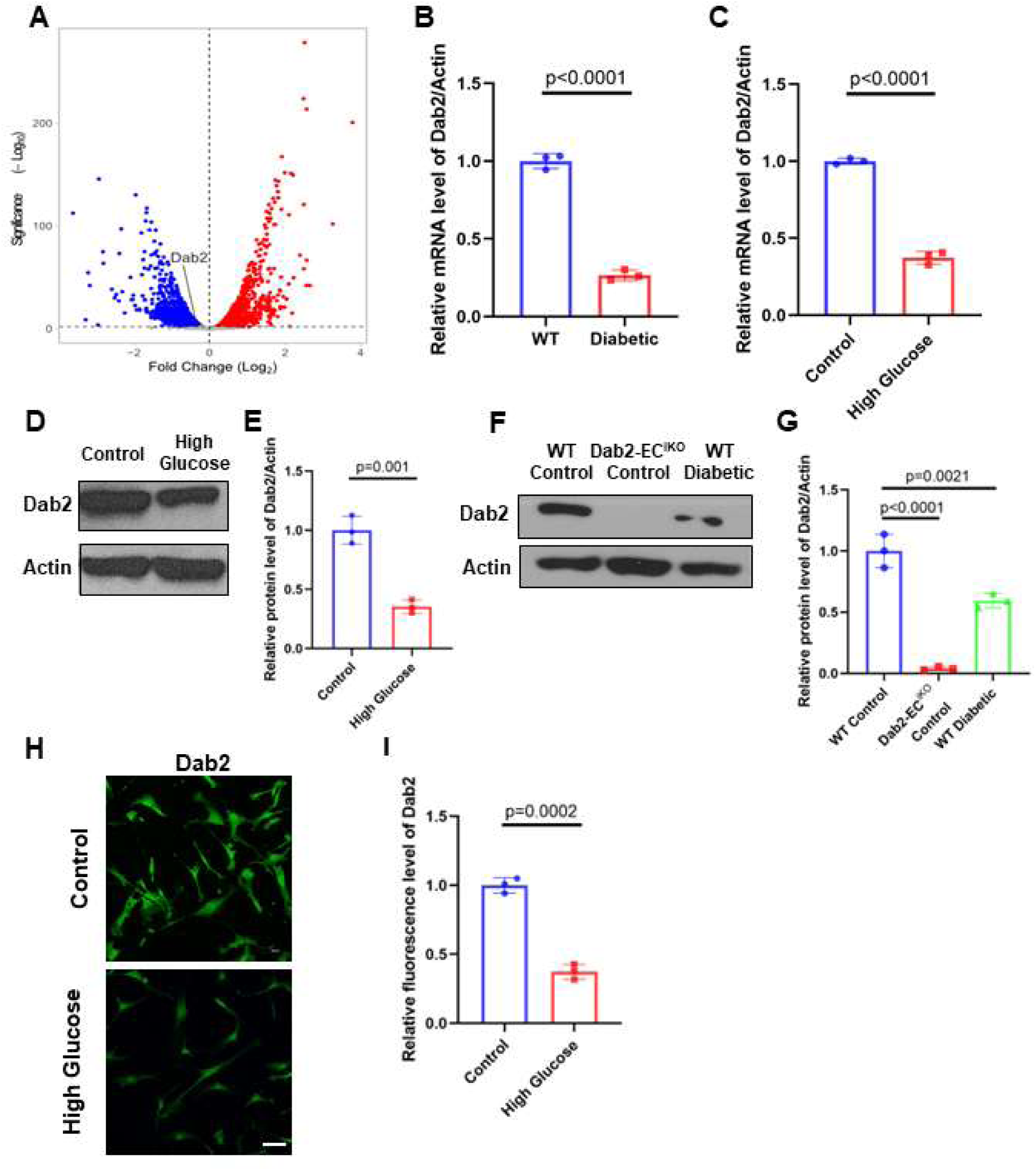
Diabetes and high glucose treatment in ECs leads to the downregulation of Dab2. (A) Volcano plot of differentially expressed genes and in skin ECs cultured in high vs normal concentration of glucose for 48 hours. The x-axis shows the log2 fold change (log2FC) and the y-axis represents the negative logarithm of the p-value (-log10 p-value) (n = 3). (B) RNA abundance of Dab2 in ECs isolated from normal or diabetic mice skin determined by qRT-PCR. (n = 3, results are presented as mean ± SD, p value calculated by t-test). (C) RNA abundance of Dab2 in skin ECs cultured in normal or high concentration of glucose determined by qRT-PCR. (n = 3, results are presented as mean ± SD, p value calculated by t-test). (D) Representative western blots of Dab2 in skin ECs cultured in normal (Control) or high concentration of glucose (E) Quantitation of protein level of Dab2 relative to Actin in (C). (n = 3, results are presented as mean ± SD, p value calculated by t-test). (F) Representative western blots of Dab2 in skin ECs isolated from WT control mice, Dab2-EC^iKO^ control mice and diabetic mice. (G) Quantitation of protein level of Dab2 relative to Actin in (E). (n = 3, results are presented as mean ± SD, p value calculated by ANOVA). (H) Representative immunofluorescence staining of Dab2 (green) in ECs treated with high or normal concentration of glucose for 24 hours. Scale bar=50 μm (I) Quantitation of fluorescence intensity in (H). (n = 3, results are presented as mean ± SD, p value calculated by t-test).

GSEA identified multiple downregulated genes in the high-glucose treatment group that are involved in the regulation of cell cycle progression such as E2F target, G2-M checkpoint, and mitotic spindle formation genes. Additionally, genes involved in cell growth, such as components of mTORC1 (mechanistic target of rapamycin complex 1) signaling and Myc target genes were downregulated (Fig. S1B-S1D). Consistent with the downregulation of Myc target genes, a majority of metabolic genes and genes involved in metabolic processes, including glycolysis and oxidative phosphorylation, were downregulated in ECs treated with a high concentration of glucose (Fig. S1B). This suggests that aerobic glycolysis, a process usually coupled with cell proliferation and growth, is impaired in the presence of a high concentration of glucose. Consistent with the RNA results, western blot analysis revealed significantly downregulated Dab2 protein levels in the diabetic group or high glucose treated group compared to non-diabetic controls (Fig. 1D-G). Downregulation of Dab2 expression in diabetic conditions was also confirmed by immunostaining of skin ECs cultured in normal or high glucose concentrations (Fig. 1H-I). Together, these *in vitro* observations suggest that hyperglycemic ECs exhibit downregulation of Dab2 mRNA and protein levels, along with reduced expression of genes involved in cellular metabolism, growth, and proliferation.

### EC-specific Dab2 knockout cause reduced angiogenesis *in vivo*

To test Dab2 function in diabetic mice we utilized a Matrigel transplantation technique where VEGFA-infused Matrigel was implanted into both wild-type (WT) and diabetic mice to create a conducive setting for *in vivo* angiogenesis analysis. This approach revealed that in the diabetic mice, the angiogenic blood vessels within the VEGFA-infused Matrigel exhibited a notable decrease in Dab2 levels compared to those in the WT mice, indicating the impact of diabetes on Dab2 expression and its potential role in angiogenesis (Fig. 2A-C). To determine if the pro-angiogenic effects of Dab2 were associated with enhanced wound healing responses *in vivo*, we examined the effects of EC-specific Dab2 deficiency on wound healing under physiological and diabetic conditions (Fig. 2C). Mice were treated with STZ and fed a high-fat diet to prepare the diabetic mouse model. Diabetes is induced by an intraperitoneal injection of STZ, with low-dose insulin given subcutaneously to manage mortality^38^. STZ selectively damages insulin-producing beta cells in the pancreas, leading to reduced insulin secretion and hyperglycemia. When used alongside an HFD, which induces insulin resistance by causing obesity, this method effectively simulates the metabolic conditions of Type 2 diabetes, eliciting both insulin resistance and compromised insulin production; thereby, creating a comprehensive diabetic mouse model.

**Figure 2.**
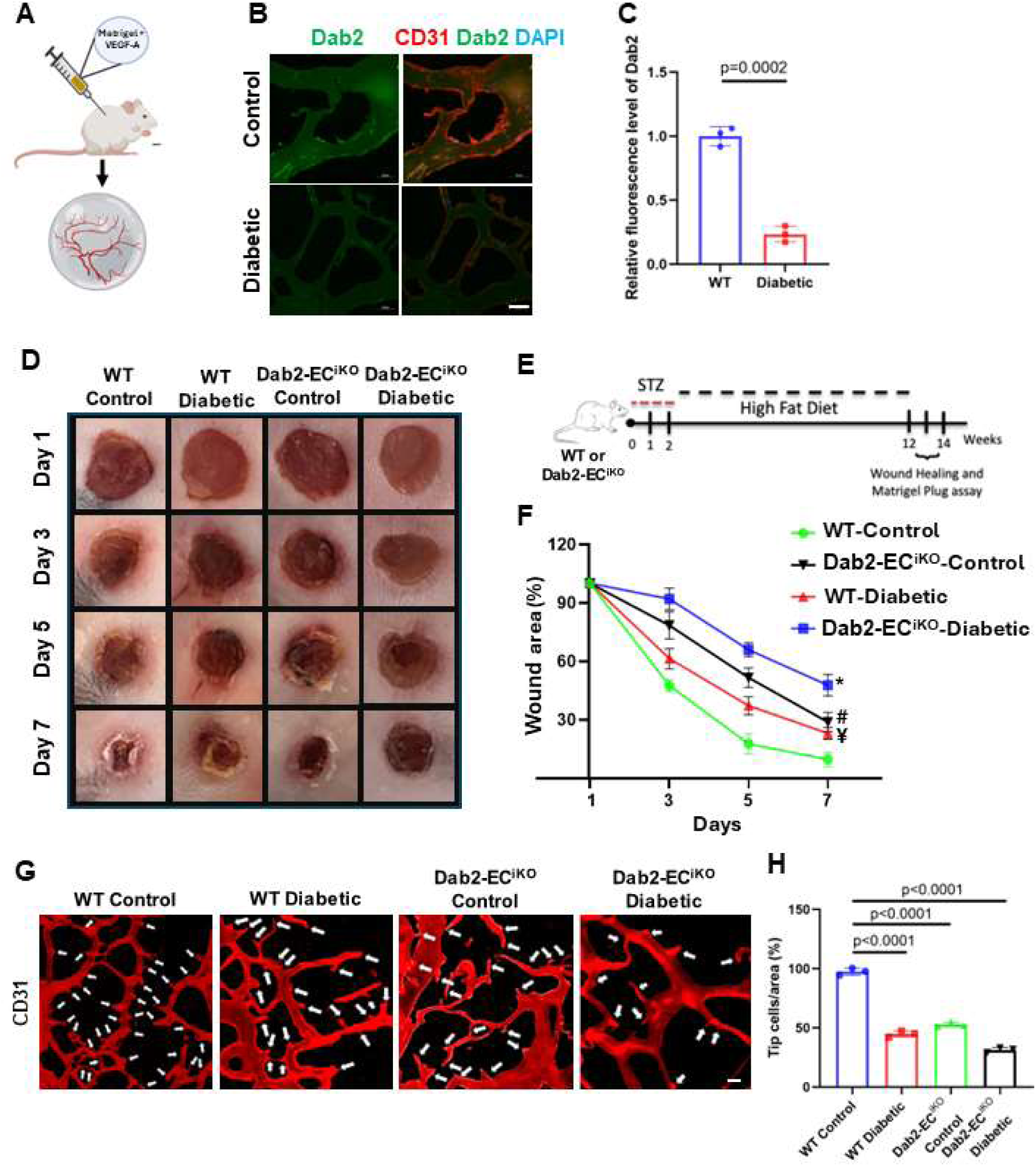
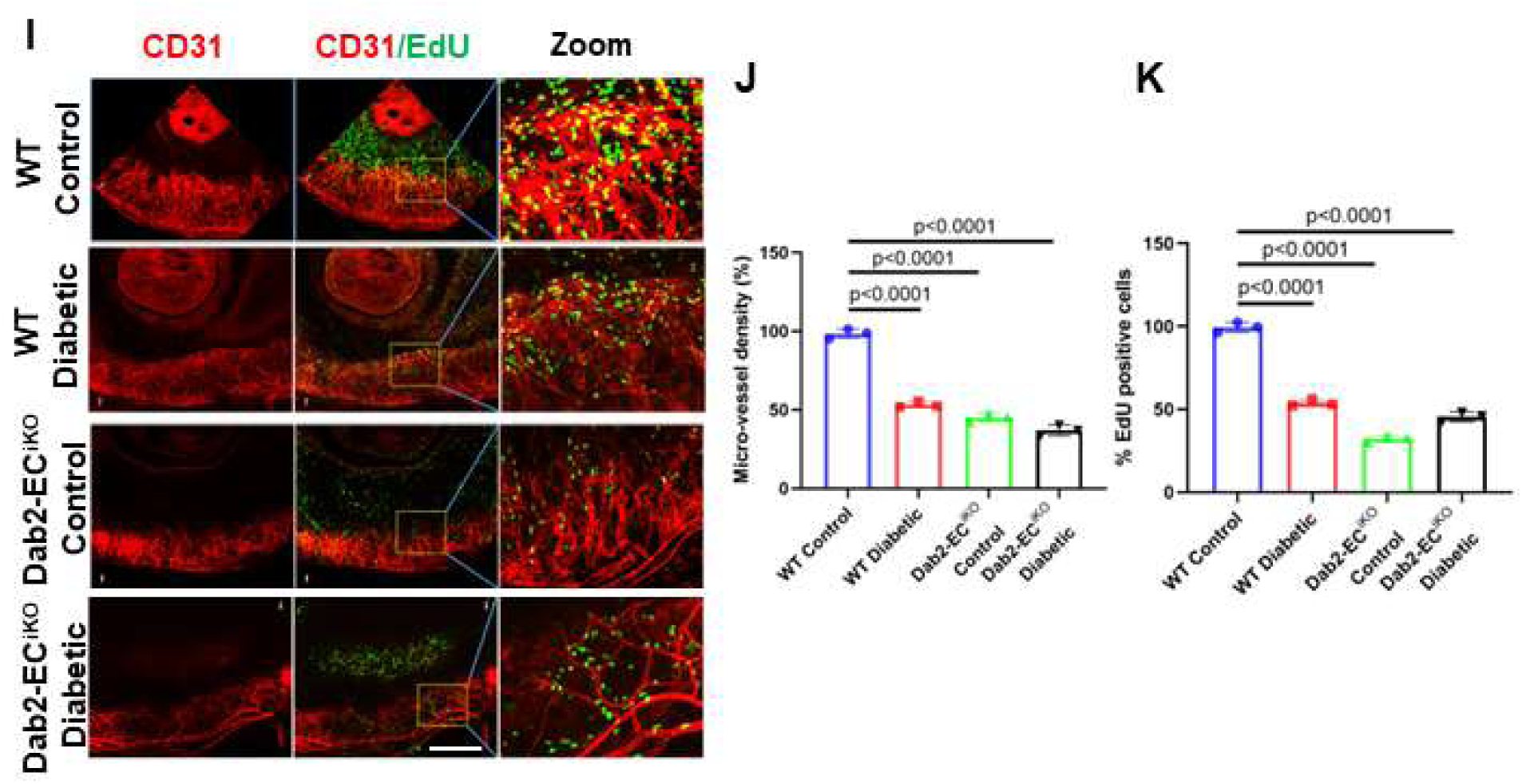
EC-specific Dab2 knockout cause reduced angiogenesis *in vivo*. (A) Schematic diagram of the Matrigel plug assay (B) Representative immunofluorescent staining of Dab2 (green) and CD31(red) in blood vessels in sections of Matrigel implant from WT and diabetic mice one week after injection. Scale bar=50μm. (C) Quantitation of Dab2 fluorescence intensity in (B). (n = 3, results are presented as mean ± SD, p value calculated by t-test). (D) Representative figures of wounds from wound healing assays in WT control mice, WT diabetes mice, Dab2-EC^iKO^ control mice and Dab2-EC^iKO^ diabetes mice. (E) Schematic diagram of the protocol used to induce diabetes in mice for the wound healing assay that illustrates the step-by-step treatment process, starting with the administration of STZ followed by a high-fat diet regimen. (F) Analysis of wound closure conducted at 1, 3, 5, and 7 days after the initial wound creation, providing a timeline view of the healing process (*P < 0.05 vs. WT mice; ^#^P<0.05 vs. WT mice; ^¥^P < 0.05 vs. WT mice. n = 6, results are presented as mean ± SD, p value calculated by ANOVA) (G) Representative immunofluorescence staining of cryosections of Matrigel plugs. Scale bar=50μm. (H) Quantitation of CD31-positive tip cell percentage in (F). (n = 3, results are presented as mean ± SD, p value calculated by ANOVA). (I) Retinal micropocket assay to assess the effect of diabetes and Dab2 deletion on angiogenesis. Scale bar=500μm. (J) Quantification of the density of the blood vessels. (n = 3, results are presented as mean ± SD, p value calculated by ANOVA). (K) Quantification of the density of EdU-positive proliferative cells. (n = 3, results are presented as mean ± SD, p value calculated by ANOVA).

**Figure 3.**
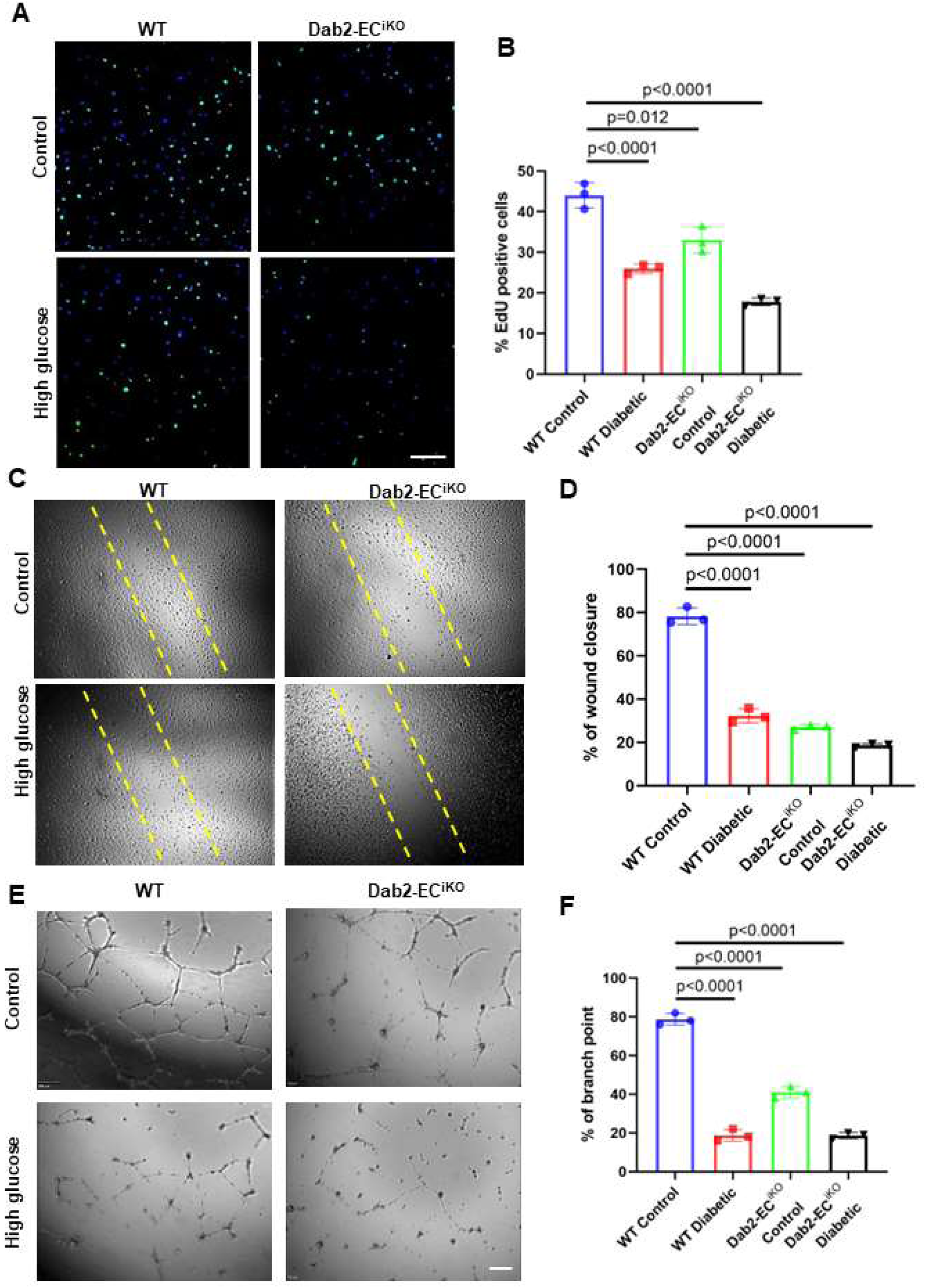
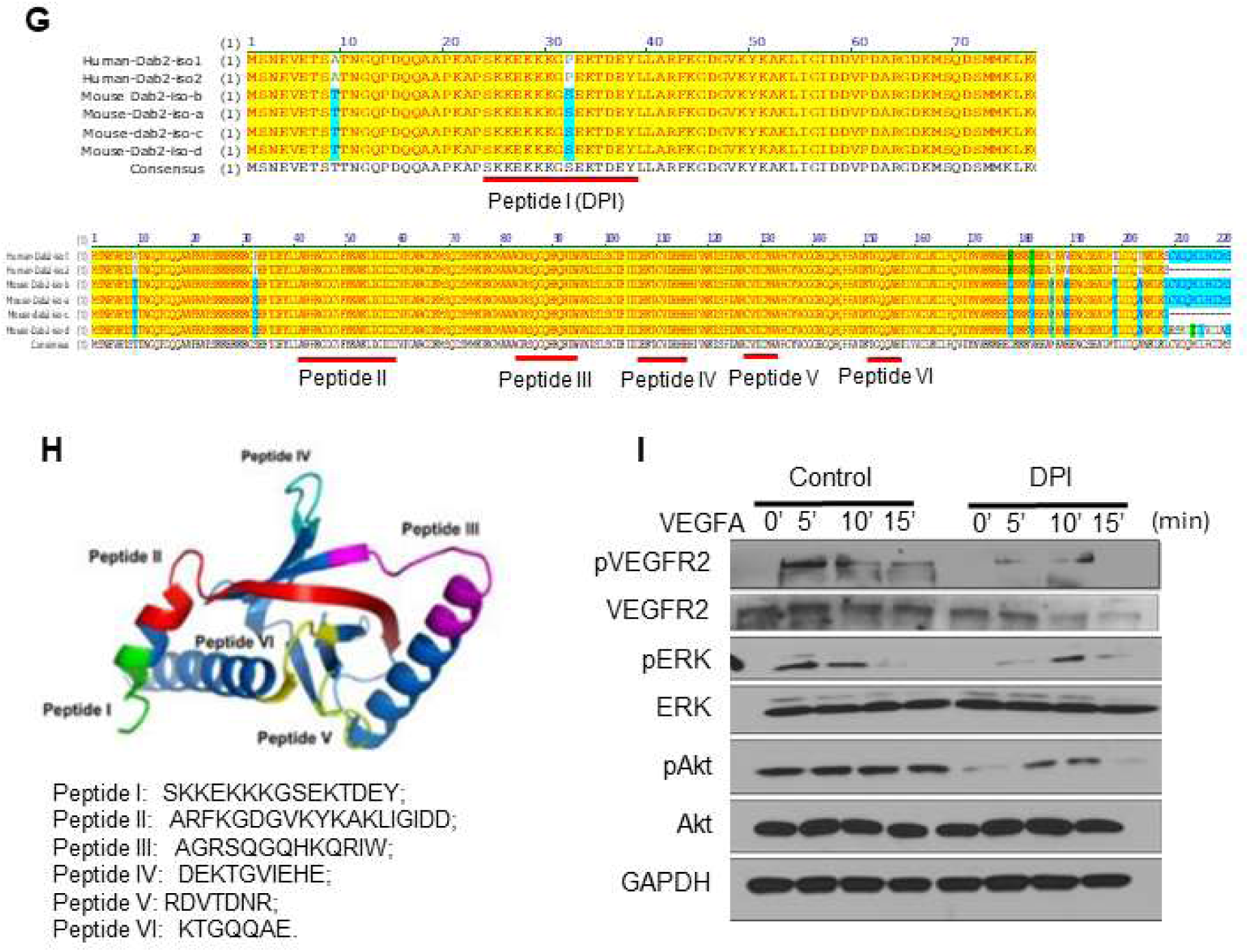
EC-specific Dab2 knockout cause reduced angiogenesis *in vitro*. (A) Representative figures of EdU incorporation (green) in WT skin ECs cultured in normal or high concentration of glucose and skin ECs from Dab2-EC^iKO^ mice with or without high concentration of glucose. Scale bar=200μm (B) Quantitation of the proportion of EdU-positive cells in (A). (n = 3, results are presented as mean ± SD, p value calculated by ANOVA). (C) Representative figures of wound closure scratch assay of ECs monolayers as described in (A). (D) Quantitation of wound closure results in (C). (n = 3, results are presented as mean ± SD, p value calculated by ANOVA). (E) Representative figures of tube formation assay of cells as described in (A). (F) Quantitation of branch points in results from (E). (n = 3, results are presented as mean ± SD, p value calculated by ANOVA). (G) Alignment of human, rat, and mouse Dab1 and Dab2 protein-coding sequences, identifying a consistent RGD motif and an additional KGD motif in the Dab2 PTB domain, suggesting evolutionarily conserved integrin binding capabilities. (H) The presence of an RGD peptide motif and an additional KGD motif in the Dab2 PTB domain suggests integrin binding capabilities. (I) Immunoblot of VEGFR2-proximal signaling components in skin ECs pretreated with DPI followed by VEGFA stimulation.

Glucose and insulin tolerance tests (ITTs and GTTs) were performed in WT mice and Dab2-EC^iKO^ mice with or without STZ injection and HFD feeding. It was found that EC-specific Dab2 deficiency led to more severe insulin resistance and higher blood glucose than WT mice (Fig S2A-B). Wounds were inflicted in the dorsal skin of normal or diabetic WT or Dab2-EC^iKO^ mice with a disposable biopsy punch under sterile conditions. Each wound was photographed at the indicated times and analyzed. EC-specific Dab2-deficiency was consistently associated with delayed wound healing response compared to WT mice (Fig. 2D-F). Whereas diabetic conditions hampered wound healing in WT mice, Dab2-EC^iKO^ diabetic mice displayed the slowest wound healing rate among all the groups. Consistently, CD31-specific immunofluorescence staining of wounds isolated on day 7 post-wound creation revealed significantly less vascularization of the wound area in diabetic and Dab2-EC^iKO^ mice (Fig. S2C-D).

To evaluate the effect of Dab2 on neovascularization *in vivo*, we subcutaneously implanted Matrigel plugs containing VEGFA into Dab2-EC^iKO^ and WT adult diabetic or control mice to directly examine EC migration and network formation *in vivo*. Consistently, the diabetic and Dab2-EC^iKO^ groups showed significantly reduced vascularization compared to WT controls (Fig. 2G-H). Furthermore, Matrigel from diabetic WT mice exhibited reduced vascularization to a similar extent to that in Dab2-EC^iKO^ mice.

To further investigate the impact of EC-specific Dab2-deficiency on angiogenesis, we performed a neo-angiogenesis assay induced by exogenous supplementation of VEGFA in the cornea using a corneal micro-pocket assay. Immunofluorescence staining of whole-mount corneas with a CD31-specific antibody confirmed the impaired vascularization and revealed reduced vessel density in Dab2-EC^iKO^ and diabetic mice (Fig. 2I-J). Specifically, we observed a significant decrease in the number of EdU-positive cells in the cornea of diabetic and Dab2-EC^iKO^ mice, indicative of a reduction in the proliferative response of ECs to VEGFA stimulation (Fig. 2K). Taken together, these observations demonstrate that EC Dab2 plays a significant role in promoting VEGFA-driven angiogenesis and wound healing *in vivo*.

### Dab2 downregulation in ECs cause reduced angiogenesis *in vitro*

To further investigate the role of Dab2 in angiogenesis, we isolated skin ECs from WT and Dab2-EC^iKO^ mice and treated them with tamoxifen. We performed *in vitro* proliferation (EdU labeling), scratch-wound healing, and EC tube formation in Matrigel in normal or high glucose media in the presence of VEGFA. The EdU-positive cells were significantly decreased in Dab2-deficient ECs compared to WT controls. Moreover, high glucose treatment led to a reduced number of EdU-positive cells in WT ECs compared to that cultured in the media with normal glucose concentration (Fig. 3A-B). Likewise, the *in vitro* scratch wound healing assay demonstrated that Dab2-deficient ECs exhibited a slower rate of wound closure compared to WT ECs. High-concentration glucose treatment of WT ECs led to blunted scratch wound closure like ECs from Dab2-EC^iKO^ mice (Fig. 3C-D). Similarly, the *in vitro* tube formation assay revealed that Dab2-deficient ECs formed fewer and less organized networks in the presence of VEGFA compared to WT ECs. (Fig. 3E-F). These findings suggest that Dab2 plays a crucial role in regulating the proliferation and migration of skin ECs under both normal and diabetic conditions. Inhibition of Dab2 expression likely underlies the compromised angiogenic function in ECs exposed to hyperglycemia in diabetes mellitus.

Dab2 is known to affect various cellular processes and is a critical regulator of VEGFR2 signaling. A previous study has shown that Dab2 could affect the VEGFR2 signaling pathway in glomerular endothelial cells^13^. However, the binding domain of Dab2 with VEGFR2 is not clear. To determine whether Dab2 could activate VEGFR2 signaling in angiogenesis and determine the precise binding domain involved in this activation, we next sought to determine exogenous inhibition of Dab2 with an inhibitory peptide (DPI) that could impede VEGFR2 signaling. We predicted a minimal peptide stretch in the Dab2 PTB domain, which is predicted to associate with VEGFR2, would abolish the interaction between VEGFR2 and Dab2 under various conditions. Using structural bioinformatics and molecular modeling, the study analyzed the alignment of human, rat, and mouse Dab1 and Dab2, identifying a consistent RGD motif and an additional KGD motif in the Dab2 PTB domain, suggesting integrin binding capabilities (Fig. 3G-H). Wild type mouse endothelial cells were pre-treated with DPI or control peptides for 18 h, followed by VEGFA stimulation. Consistent with the observations made in Dab2-depleted ECs, the DPI peptide significantly reduced the levels of pVEGFR2, pAkt, and pERK relative to the control peptide (Fig. 3I). Together, these results suggest that Dab2 plays a crucial role in regulating the VEGF-VEGFR2 signaling pathway in EC angiogenesis.

### Restoration of Dab2 Expression in ECs Rescues Impaired Angiogenesis and Wound Healing in Diabetic Mice

To investigate the therapeutic efficacy of Dab2 restoration in diabetic conditions, we conducted a rescue experiment in STZ-induced diabetic mice. We intravenously administered Dab2-mRNA encapsulated in LNPs conjugated with Lyp1 peptide at a dose of 15 μg/mouse, twice a week, during the wound healing period (Fig. S3). This treatment was aimed at restoring Dab2 expression and enhancing angiogenesis and wound healing. Diabetic mice treated with LNPs-LyP1-Dab2-mRNA exhibited significantly accelerated wound healing activity compared with the untreated diabetic mice (Fig. 4A-B). The treated group exhibited accelerated wound closure and healing rates, with most wounds nearly fully healed by day 7, in contrast to the control group (LNPs-LyP1-GFP mRNA), where wound healing was significantly impeded within the same time. To complement these *in vivo* findings, we utilized a Dab2-Lentivirus system to overexpress Dab2 in high glucose-treated primary skin ECs enriched from adult mice. Dab2 overexpression in high glucose treatment recovered the proliferative capacity in ECs compared to control (Fig. 4C-D). We also performed scratch assays and tube formation assays to mimic wound healing and angiogenesis *in vitro* (Fig. 4E-H). Cells overexpressing Dab2 tended to form more organized and complex tube-like structures, as revealed by tube formation assays (Fig. 4E-F) and exhibited increased migratory behavior in scratch assays (Fig. 4G-H), compared to control cells. These results confirmed the ECs autonomous pro-angiogenic function of Dab2 and highlight the potential therapeutic utility of overexpressing Dab2 to restore impaired angiogenic responses in diabetic conditions.

**Figure 4.**
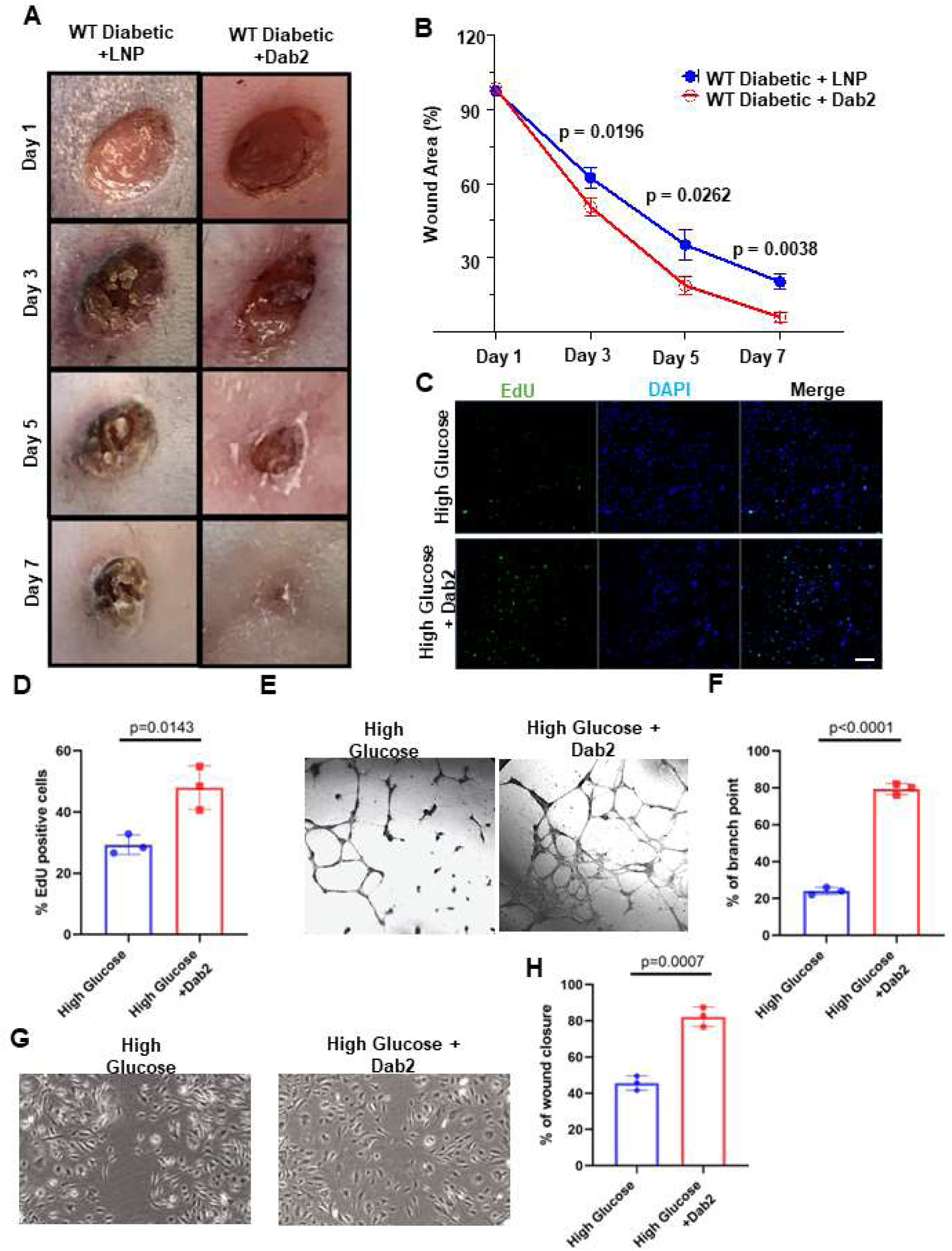
Restoration of Dab2 expression in ECs rescues impaired angiogenesis and wound healing in diabetic mice. (A) Representative figures of wounds from wound healing assays in diabetic mice treated with LNPs carrying mRNAs encoding GFP or Dab2. (B) Quantitation of wound closure conducted on day 1, 3, 5, and 7 after the initial wound (n=5, p value calculated using t-test). (C) Representative figures of EdU of skin ECs cultured in high concentration of glucose infected with empty lentivirus vector or lentivirus carrying Dab2 cDNA. Scale bar=200μm. (D) Quantitation of the proportion of EdU-positive cells in (C). (n=3, p value calculated using t-test). (E) Representative figures of tube formation assay on skin ECs cultured in high concentration of glucose and infection empty vector or lentivirus carrying Dab2 cDNA. (F) Quantitation of branch points in results from (E). (n = 3, results are presented as mean ± SD, p value calculated by t-test). (G) Representative figures of wound healing assay on skin ECs cultured in high concentration of glucose and infection empty vector or lentivirus carrying Dab2 cDNA. (H) Quantitation of wound closure results in (G). (n = 3, results are presented as mean ± SD, p value calculated by t-test).

### FoxM1 is downregulated in diabetes and regulates Dab2 transcription

To find the mechanism of Dab2 regulation, we further explored the RNA sequencing data in Figure 1. Volcano plot analysis revealed downregulation of transcription factors involved in cell proliferation and growth, including FoxM1 (Forkhead box M1), Egr2 (Early growth response protein 2), Hes1 (Hairy and enhancer of split 1), Etv4 (ETS variant 4 exclusively in the high glucose-treated samples compared with the control group (Fig. 5A, S1A). qPCR analysis showed that high glucose treatment of ECs from diabetic mice resulted in the downregulation of Dab2 (Fig. 5B-C). Similar to the qPCR results, western blot analysis revealed significantly downregulated FoxM1 protein levels in the diabetic group or high glucose treated group compared to non-diabetic controls (Fig. 5D-G).

**Figure 5.**
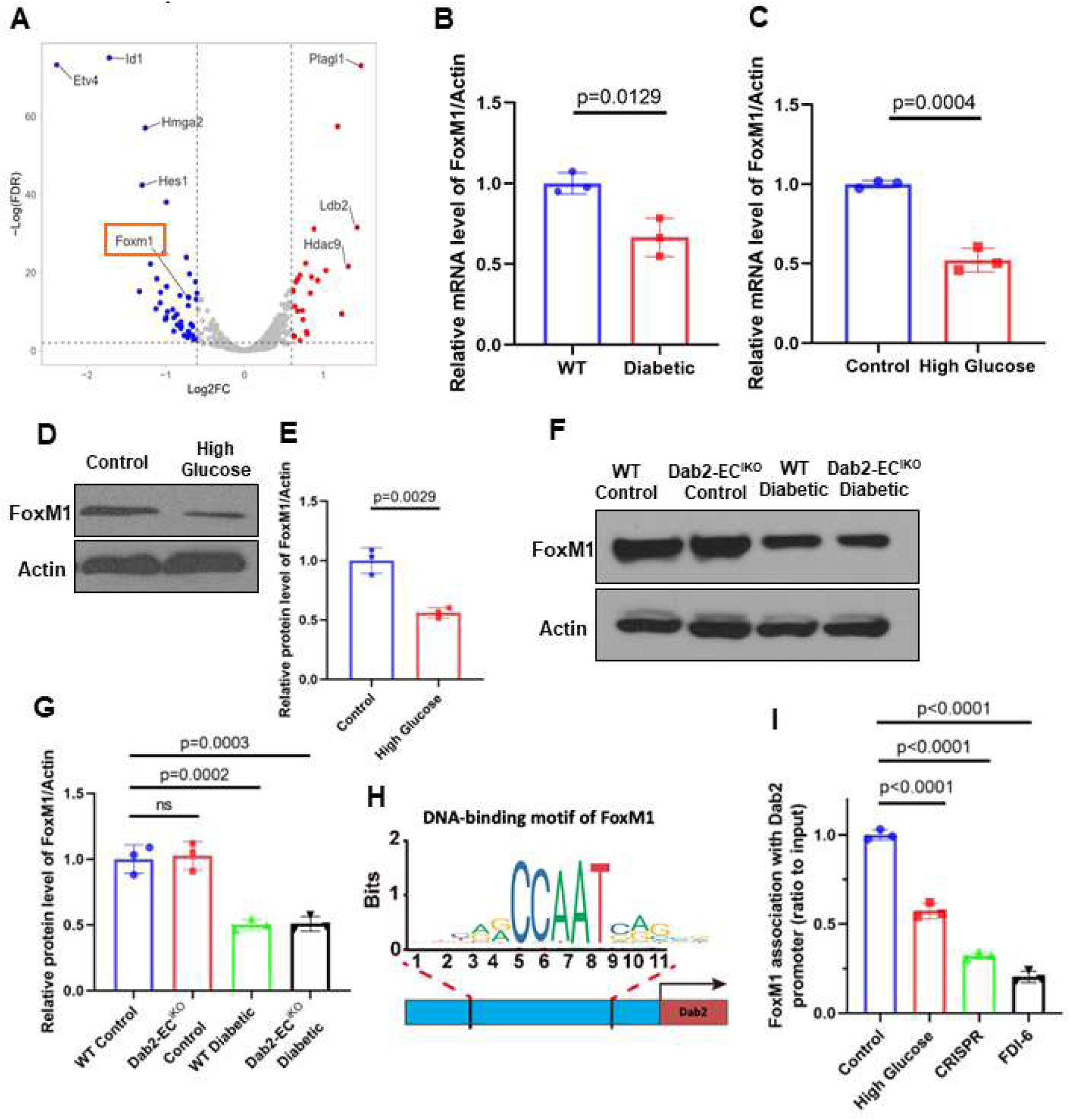
Foxm1 is downregulated in diabetes and regulates Dab2 transcription. (A) Volcano plot of differentially expressed transcription factors in skin ECs cultured in high vs normal concentration of glucose for 24 hours. The x-axis shows the log2 fold change (log2FC) and the y-axis represents the negative logarithm of the p-value (-log10 p-value). n = 3. (B) RNA abundance of Foxm1 in ECs isolated from normal or diabetic mice skin determined by qRT-PCR. (n = 3, results are presented as mean ± SD, p value calculated by t-test). (C) RNA abundance of Foxm1 in skin ECs cultured in normal or high concentration of glucose determined by qRT-PCR. (n = 3, results are presented as mean ± SD, p value calculated by t-test). (D) Representative western blots of Dab2 in skin ECs cultured in control or high concentration of glucose. (E) Quantitation of protein level of Dab2 relative to Actin in (D). (n = 3, results are presented as mean ± SD, p value calculated by t-test). (F) Representative western blots of Dab2 in skin ECs isolated from WT control mice, Dab2-EC^iKO^ control mice, WT diabetic mice and Dab2-EC^iKO^ diabetic mice. (G) Quantitation of protein level of Dab2 relative to Actin in (E). (n = 3, results are presented as mean ± SD, p value calculated by t-test). (H) JASPAR-predicted FoxM1-Binding site in the Dab2 promoter. (I) FOXM1 binding to the Dab2 promoter in ECs exposed to high concentration of glucose, or FDI-6, or with a CRISPR-mediated deletion mutation in the FoxM1 binding site on the Dab2 promoter. (n = 3, results are presented as mean ± SD, p value calculated by ANOVA).

The concomitant downregulation of FoxM1 and Dab2 transcripts under diabetic conditions raises the possibility that, as a regulatory transcription factor, FoxM1 directly influences Dab2 expression to modulate VEGFR2-mediated endothelial function. FoxM1 has been shown to promote cell cycle progression, cell proliferation, and cellular metabolism^39,40^, while Dab2 is a critical regulator of VEGFR2 signaling. To investigate how FoxM1 regulates Dab2 expression and VEGFR2 signaling, we used ChIP-qPCR analysis in ECs to determine if FoxM1 can bind to the Dab2-promoter and other regulatory regions by using ChIP-qPCR analysis in ECs isolated from wild type mice. We used the JASPAR database^41,42^ to predict potential FoxM1 transcription factor binding sites. A potential binding site within the promoter region of the Dab2 locus with the highest score was selected for further study (Fig. 5H, S4A). ChIP-qPCR results showed that indeed FoxM1 binds to this predicted region within the promoter region of Dab2 in ECs. Interestingly, the binding of FoxM1 to the Dab2 promoter region is decreased when the ECs were treated with high-concentration glucose or the FoxM1 inhibitor FDI-6 (Fig. 5I). More importantly, disrupting the FoxM1 binding site with CRISPR/Cas-induced mutation in ECs also significantly diminished FoxM1 binding. Our data demonstrate that FoxM1 directly binds to the promoter region of Dab2 and disruption of this binding regulates its transcription.

### FoxM1 inhibitor FDI-6 downregulates Dab2 expression and the phosphorylation of VEGFA-induced VEGFR2

The significance of FoxM1 in ECs and vascular repair is underscored by its role in promoting endothelial regeneration and resolving inflammatory lung injury. FoxM1, expressed during embryogenesis in various cell types, including ECs, is crucial for pulmonary vascular ECs proliferation and endothelial barrier recovery post-inflammatory injury. Notably, FoxM1 facilitates the reannealing of endothelial adheres junctions, enhancing endothelial barrier function post-vascular injury. Aging-related impairment in endothelial regeneration and inflammatory injury resolution is linked to inadequate FoxM1 induction, which, when addressed through transgenic expression, improves outcomes in aged mice, highlighting the potential of FoxM1 as a target for vascular repair interventions^43–46^.

To determine if FoxM1 controls Dab2 transcription, we used the small molecule FDI-6 to specifically inhibit FoxM1 function. FDI-6 is a FoxM1 transactivational inhibitor that blocks its DNA binding^47,48^. Immunofluorescence staining of primary ECs revealed a significant decrease in the expression of pVEGFR2 and Dab2 in FDI-6-treated conditions compared to the control group, while the expression of FoxM1 remained unchanged (Fig 6A-C). In the VEGF signaling pathway, ERK and AKT would normally be phosphorylated and activated. In high glucose treatments, we observed diminished phosphorylation and activation of VEGFR2 (pVEGFR2), Akt (pAkt), and ERK (pERK) (Fig. 6D-E). Furthermore, FDI-6 treatment reduced the protein levels of pVEGFR2 as well as activation of both pERK and pAKT (Fig. 6F-G). In contrast, the total protein level of FoxM1 was unaffected by FDI-6 treatment. These observations suggest that FoxM1 regulates the expression of Dab2 which, in turn, controls VEGFR2 signaling.

**Figure 6.**
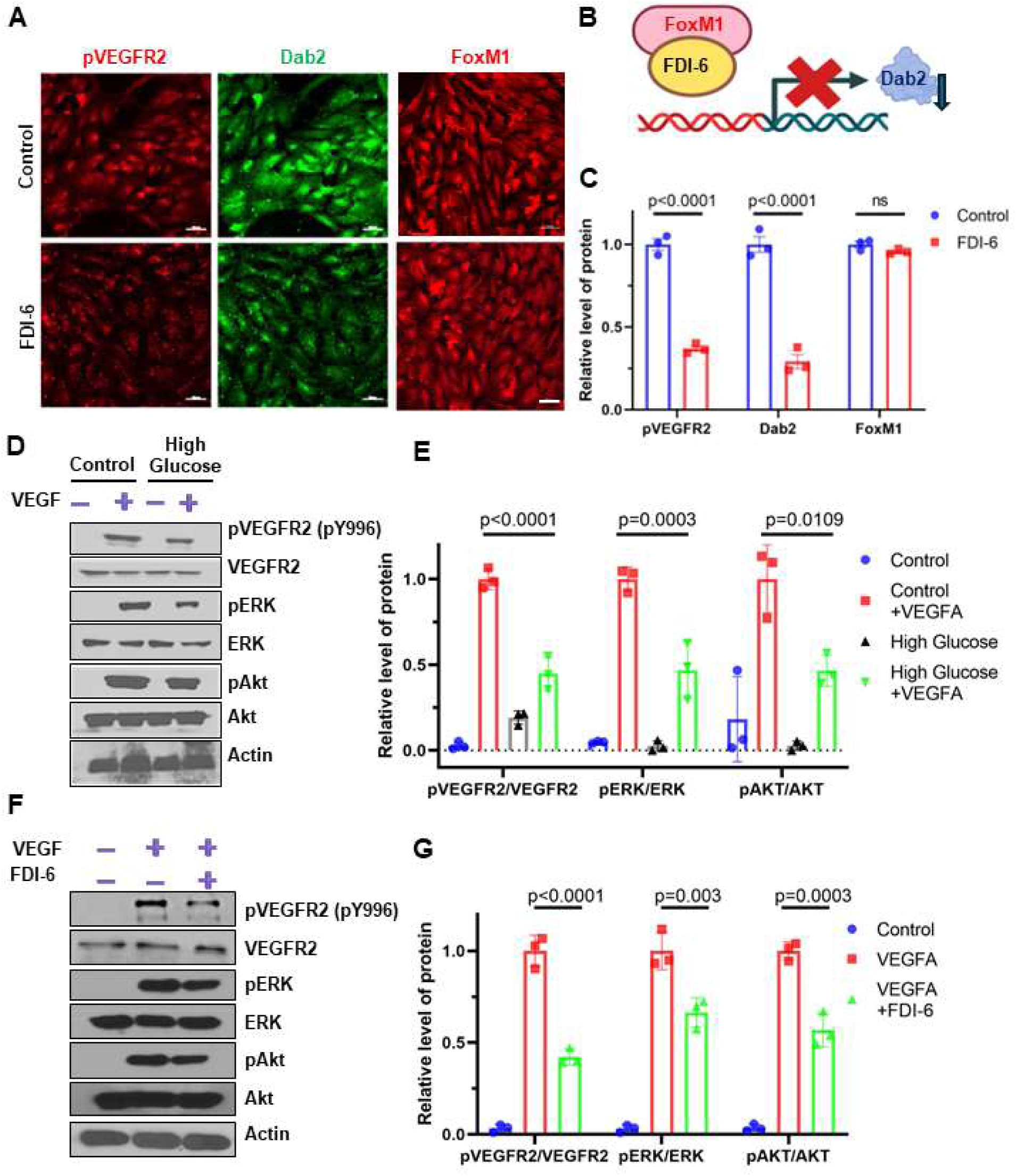
Foxm1 inhibitor FDI-6 downregulates Dab2 expression and the phosphorylation of VEGFA-induced VEGFR2. (A) Representative immunofluorescence staining of skin ECs treated with or without FDI-6. Scale bar=50 μm. (B) Schematic diagram showing inhibition of Dab2 expression by FDI-6 (C) Quantitation of the immunofluorescence intensity in (A). (n = 3, results are presented as mean ± SD, p value calculated by t-test). (D) VEGFA-induced phosphorylation of key VEGFR2-proximal signaling components in skin ECs treated in control or high glucose concentration with or without VEGF assessed by Western blot analysis. (E) Quantitation of results described in (D). (n = 3, results are presented as mean ± SD, p value calculated by ANOVA). (F) Representative of Western blot of VEGFA-induced key VEGFR2-proximal signaling in skin ECs treated with or without FDI-6. (G) Quantitative analysis of immunoblots in (C). (n = 3, results are presented as mean ± SD, p value calculated by ANOVA).

## Discussion

Chronic non-healing wounds in diabetes present a complex challenge and impaired angiogenesis appears to underpin these defects in tissue repair and regeneration. Blunted angiogenesis is often attributed to dysregulated signaling and aberrant gene expression precipitated by diabetic conditions. Therefore, identifying crucial angiogenesis modulators is crucial to illustrate the pathogenesis of blunted angiogenesis in diabetes and assess their therapeutic utility. Our observations here provide important insights into the precise involvement of the highly conserved endocytic adaptor protein, Dab2, and its transcriptional regulation during wound-induced angiogenesis in diabetes. Dab2 is widely expressed in various tissues, suggesting tissue-specific and context-dependent roles in multiple physiological processes^49^. Dab2 primarily functions as a cytosolic, clathrin and cargo binding adaptor protein, with a pivotal role in endocytosis. By facilitating the internalization of cargo molecules and the binding of clathrin-coated vesicles to cargo proteins through clathrin-mediated endocytosis, Dab2 not only facilitates signal transduction and receptor recycling but also plays a crucial role in maintaining cellular homeostasis and regulating intracellular trafficking^50,51^.

Intriguingly, Dab2 and its phosphorylated form are enriched in ECs^13,52^. Studies in *Xenopus* and *Zebrafish* have revealed a functional contribution of Dab2 to developmental angiogenesis through both VEGF-dependent and VEGF-independent mechanisms^53–56^. Earlier *in vitro* studies revealed that Dab2 function is conducive to EC migration via mitigating VEGF signaling^49,57^. Similarly, Dab2 is required for the vascularization of the brain tissue and the establishment of the neurovascular unit in part by enhancing VEGF signaling^58,59^. At least two other studies suggested that activated receptor endocytosis through Dab2 results in augmented VEGF signaling in ECs^14,56^. Nakayama *et al.* showed that phosphorylation of its phosphotyrosine-binding (PTB) domain diminishes its interaction with the VEGF pathway receptors, VEGFR-2 and VEGFR-3, revealing the specificity of Dab2 for VEGFRs^14^.However, the upstream regulatory mechanisms moderating the expression of Dab2 and its interaction with VEGFR2 remain unknown. Furthermore, despite the well-established pro-angiogenic role of Dab2, the therapeutic potential of exploiting Dab2 mediated angiogenesis to enhance wound healing in a disease context (*i.e.*, diabetic wounds), has remained unexplored.

In our quest to investigate the involvement of endocytic adaptor proteins in mitigating aspects of diabetes, we identified Dab2 via our bulk RNA-sequencing analysis of hyperglycemic ECs as one of the downregulated genes. Similarly, we observed that ECs isolated from the STZ-induced diabetic mice model, showed a significant downregulation of Dab2 mRNA and protein levels. Although the precise involvement of Dab2 in diabetes remains to be addressed, Dab2 appears to be involved in regulating blood glucose metabolism and its deficiency at least in myeloid cells, has been implicated in compromised glucose tolerance in mice^60^. More intriguingly, and only recently, polymorphisms in the Dab2 gene have been associated with type 2 Diabetes Mellitus (T2DM) in a recent population-based study ^55^.

Because of the lack of information about the precise role of dab2 in diabetes, we went on to develop an EC-specific Dab2-deficient mouse model to examine the effects of Dab2 deletion on angiogenesis in diabetic conditions. we observed that endothelial-specific loss Dab2 was strongly associated with delayed wound healing response and diabetic background further exacerbated this process in Dab2-EC^iKO^ mice, coupled with severely blunted angiogenesis. Meanwhile our *in vitro* model revealed that loss of endothelial Dab2 resulted in curtailed angiogenesis, as evidenced by diminished cell migration, network formation, and proliferation.

Mechanistically, Dab2 deletion affected VEGFR2 activation, thereby impacting vascular development both *in vitro* and *in vivo*. While the DPI, which blocked the certain domain that Dab2 interact with VEGFR2, could downregulate the VEGF2 activation in VEGFA stimulated ECs. These results align with prior studies on the role of Dab2 in angiogenesis but introduce novel insights, including a comprehensive *in vivo* evaluation of angiogenesis in the diabetic context. This unique approach enhances our understanding in dissecting the pathological mechanisms of Dab2 underlying diabetic vascular complications.

Delayed wound healing in diabetic patients presents a significant clinical challenge while the inhibitory effect of endothelial-specific Dab2 knockout on wound healing in our result suggested that exogenous supplementation of Dab2 could be a potential therapeutic approach. Consequently, we investigated the effects of exogenous Dab2 restoration on angiogenesis *in vivo* and *in vitro*. In this study, we employed a novel method to administer Dab2 supplementation via the deployment of *Dab2*-mRNA encapsulated in LNP conjugated with Lyp1 peptide. This novel therapeutic approach bypasses common encumbrances associated with other modes of delivery, such as inadequate absorption or shorter half-life of delivered proteins. Consistently, a recent study has demonstrated the safety and specificity of augmenting VEGFA via nanoparticles^61^. The LNPs-Lyp1-Dab2 mRNA group exhibited significantly accelerated wound healing activity compared with the untreated diabetic siblings, as observed by enhanced wound closure and healing rates, with most wounds fully healed by day 7. This result demonstrated the feasibility and positive effect of Dab2 mRNA supplementation *in vivo*. The beneficial effects of Dab2 restoration on angiogenesis also suggested that supplementation of Dab2 may serve as a possible therapeutic target for diabetes patients delayed wound healing.

Further delving into the mechanism, we demonstrated that FoxM1, a highly conserved transcription factor, exerts regulatory control over Dab2 expression in ECs during angiogenesis associated with wound healing. Notably, we showed that FoxM1 binds to the Dab2 promoter to drive Dab2 expression. Therefore, we have established that FoxM1 exerts a positive regulatory control on Dab2 transcription, adding further insights into the molecular mechanisms regulating angiogenesis during wound healing in diabetic conditions.

Although previous studies have suggested an involvement of FoxM1 in the pathogenesis of diabetes, emerging studies have revealed that deletion of FoxM1 in diabetic mice impairs wound healing, in part, via impeded recruitment of immune cells^62^. However, the endothelial-specific role of FoxM1 in diabetic wounds was not addressed. FoxM1 plays a crucial role in β-cell proliferation, essential for pancreatic repair and insulin secretion^63^. It activates pathways critical for β-cell growth and interacts with regulatory genes, enhancing its transcriptional activity via the insulin receptor-mediated pathway. Additionally, FoxM1’s involvement extends to nutrition-induced β-cell growth and its significance in gestational diabetes, highlighting its importance in β-cell function across various physiological conditions^61,63^. While previous studies acknowledge the pivotal role of FoxM1 in diabetes-related cellular functions and β-cell proliferation, our study delves into the mechanistic interaction between FoxM1 and Dab2 within the context of ECs and angiogenesis during wound healing in diabetic conditions.

Our study provides crucial insights into the role of Dab2 and FoxM1 in diabetic wound healing, highlighting the therapeutic potential of Dab2 mRNA encapsulated in lipid nanoparticles. This novel approach not only advances our understanding of the molecular mechanisms underlying diabetes-related angiogenesis and wound repair but also opens new avenues for developing targeted treatments for diabetic complications.

## Acknowledgments

S.B. and H.C. conceived the project.

S.B., Y.W.L., S.E.B., H.W, A.E.B., J.G., S.W., Y.S., J.X., J.S., H.C. participated in experimental design, execution, and data analysis.

S.B., A.E.B. contributed to Mouse corneal micropocket assay.

S.B., Y.W.L., H.C. analyzed the data of bulk RNA-seq.

S.B., Y.W.L., designed and performed CRISPR/Cas9 experiment.

S.B., H.W., performed G.T.T and I.T.T assays.

Y.S., J.S., designed and prepared the nanoparticles.

S.B. prepared lentivirus.

S.B. performed immunohistochemistry and confocal imaging studies.

S.B., H.W., performed Western blot analysis.

S.B., H.W., performed the *in vitro* and *in vivo* angiogenesis and wound healing assay.

S.B., S.E.B., did the quantification and statistics analysis.

S.B., S.E.B., J.G., D.B.C. and H.C. wrote the manuscript.

S.B., S.E.B., H.W., J.G., Y.W.L., K.L., W.W., D.B.C., J.X., M.A., J.S., H.C. reviewed and edited the manuscript.

---

**Figure S1.**
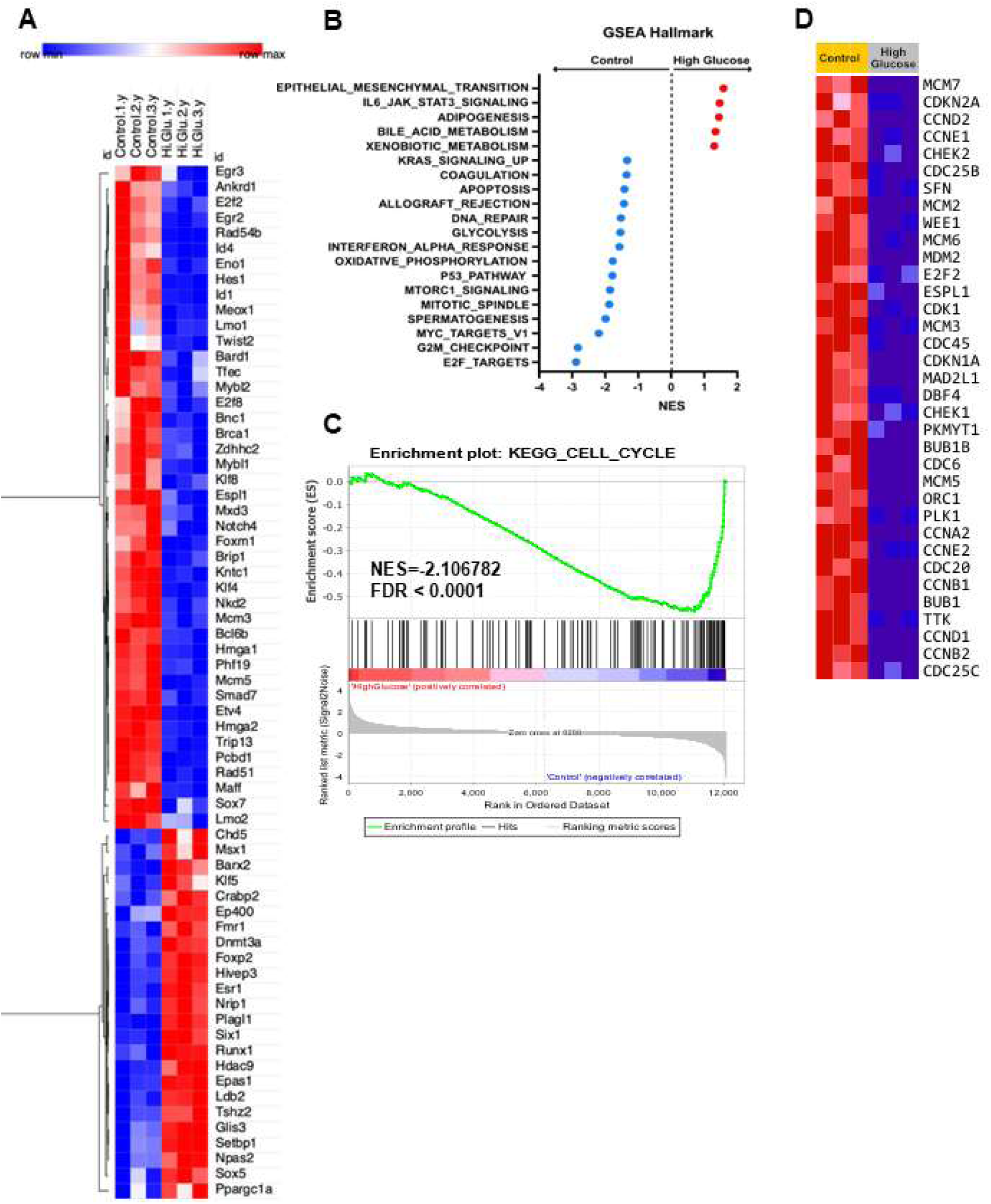
Heatmaps showing differentially regulated gene expression in CD31-enriched primary mouse skin ECs exposed to normal or high glucose concentrations for 48 hours. (A) Differential gene expression analysis revealed 168 significantly downregulated and 386 significantly up-regulated genes in the ECs grown in high glucose culture conditions compared to ECs cultured in normal glucose media. Sample genes are shown (n = 3). (B) GSEA Hallmark genesets enriched on differentially expressed genes from bulk RNA-seq data described in bulk RNA-sequencing. Geneset with a nominal p-value <0.05 and FDR <0.25 were included. Normalized Enrichment Score (NES) for all gene sets are shown. (C) Enrichment plot for KEGG Cell Cycle. (D) Multiple down-regulated genes in the high-glucose treatment group are involved in the regulation of cell cycle progression (n = 3).

**Figure S2.**
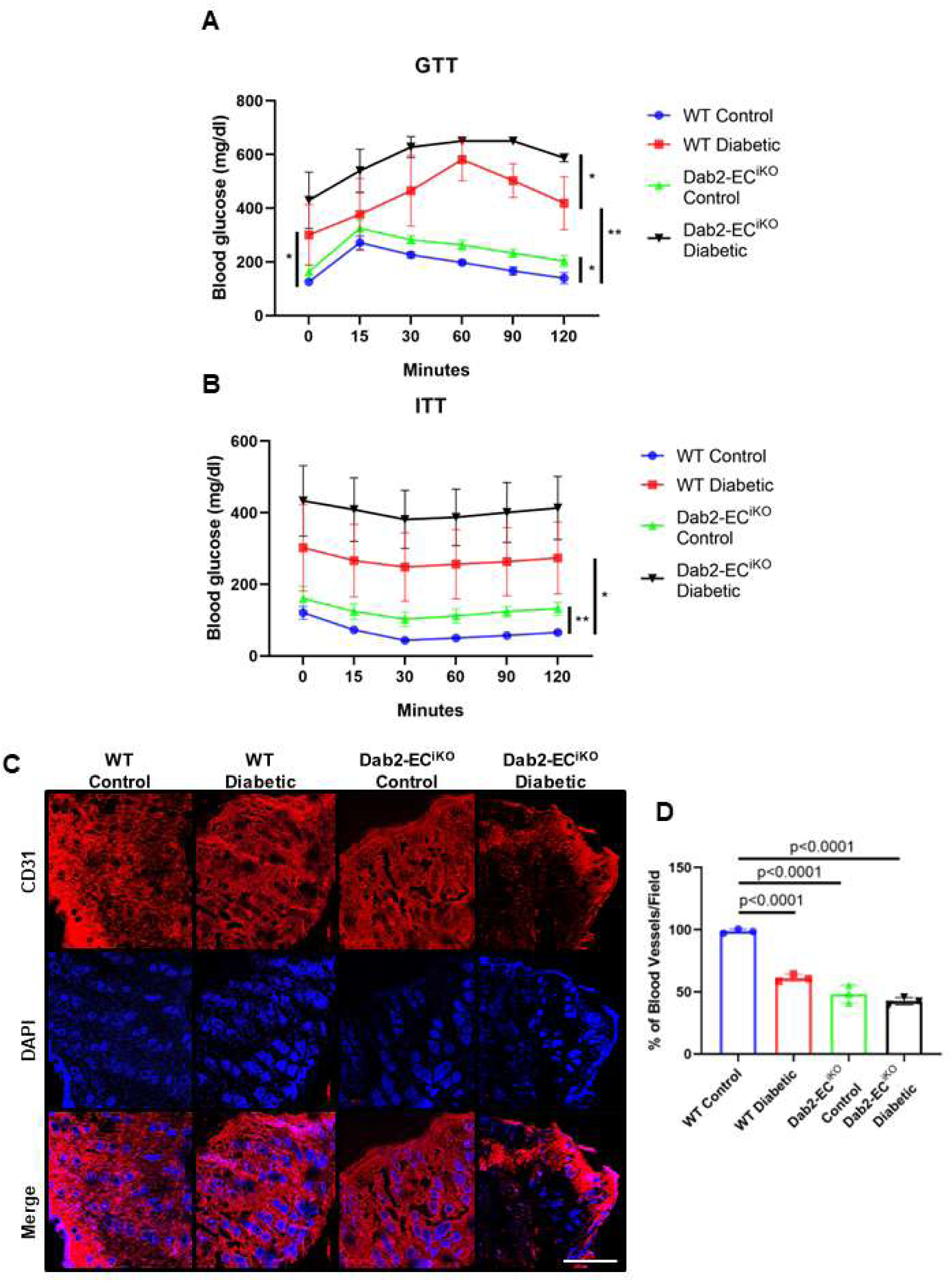
GTT and ITT of HFD WT mice and Dab2-EC^iKO^ mice, and wound CD31 in diabetic WT mice and Dab2-EC^iKO^ mice. (A) Glucose tolerance test (GTT) of WT and Dab2-EC^iKO^ mice with or without diabetes after low-dose STZ injection and 12-weeks HFD feeding described in Figure 2E. (n = 3-6, results are presented as mean ± SD, p value calculated by Student’s t-test, *p<0.05, **p<0.01). (B) Insulin tolerance test (ITT) of WT and Dab2-EC^iKO^ mice with or without diabetes after low-dose STZ injection and 12-weeks HFD feeding described in Figure 2E. (n = 5, results are presented as mean ± SD, p value calculated by Student’s t-test, *p<0.05, **p<0.01). (C) Representative immunofluorescence staining of CD31 (red) in wound area from collected in mice described in Figure 2D. Scale bar=100μm. (D) Quantitation of CD31-positive blood vessel density in Figure 2D. (n = 3, results are presented as mean ± SD, p value calculated by ANOVA).

**Figure S3.**
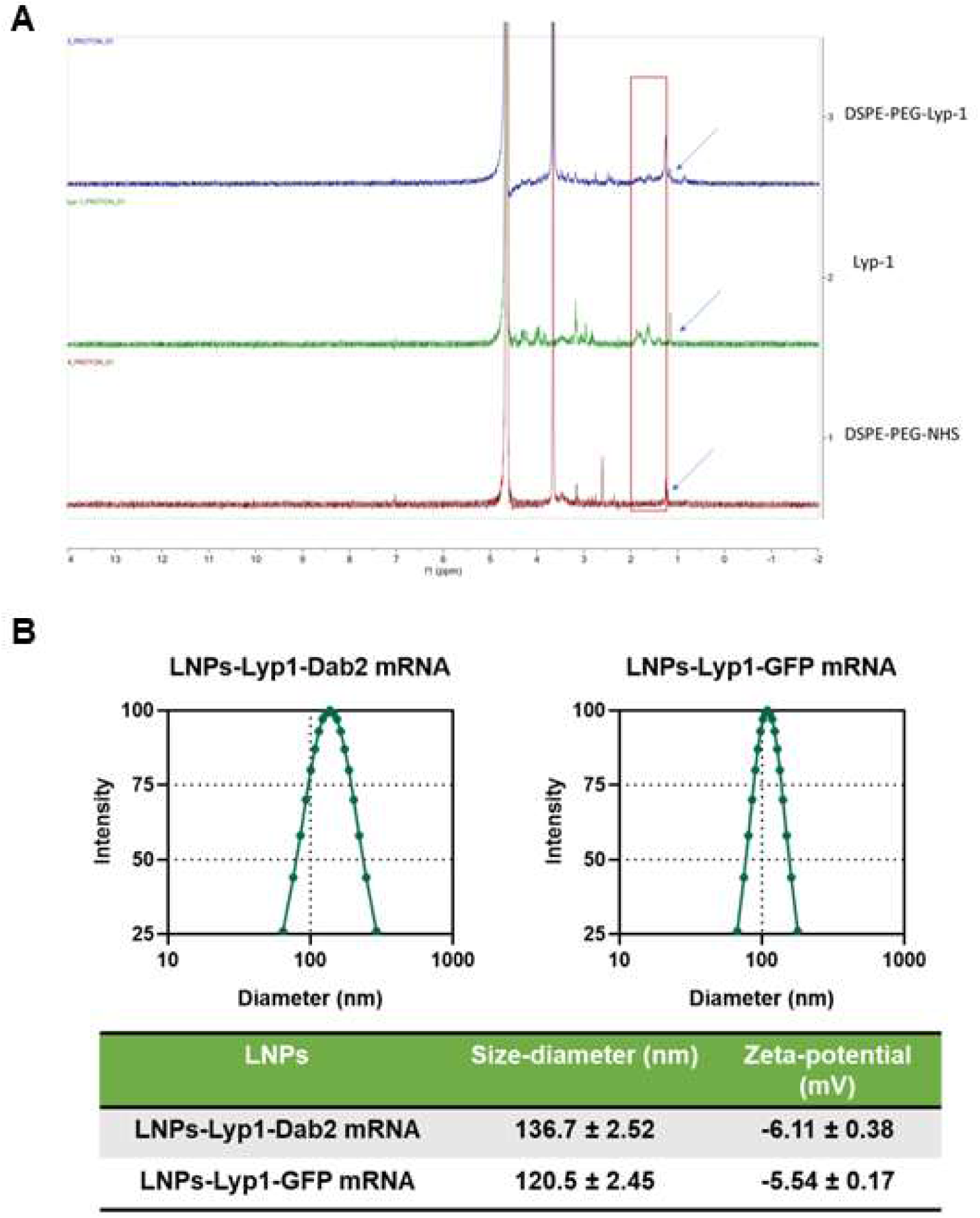
Characterization of the Lyp1-LNP-Dab2 mRNA- and Lyp1-LNP-GFP mRNA-containing lipid nanoparticles (LNPs). (A) ^1^H-NMR to characterize the successful synthesis of DSPG-PEG-Lyp1. (B) Size and Zeta potential of LNPs-Lyp1-Dab2 mRNA and LNPs-Lyp1-GFP mRNA (control group).

**Figure S4.**
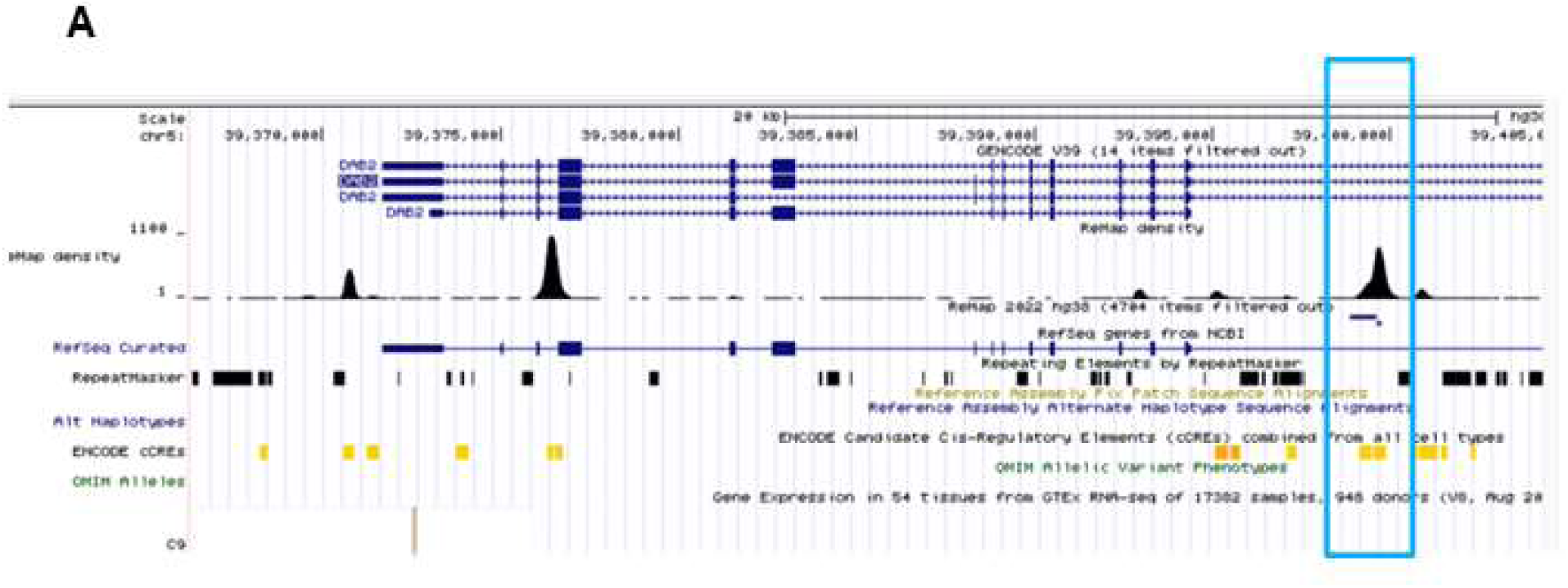
FoxM1 binding zone in Dab2 promoter. (A) UCSC genome viewer display of FoxM1 binding sites (boxed peak) in Dab2 promoter region. The x-axis represents the genomic region of the Dab2 promoter, the y-axis shows the peak height of the FoxM1 binding sites.

**Table S1.**
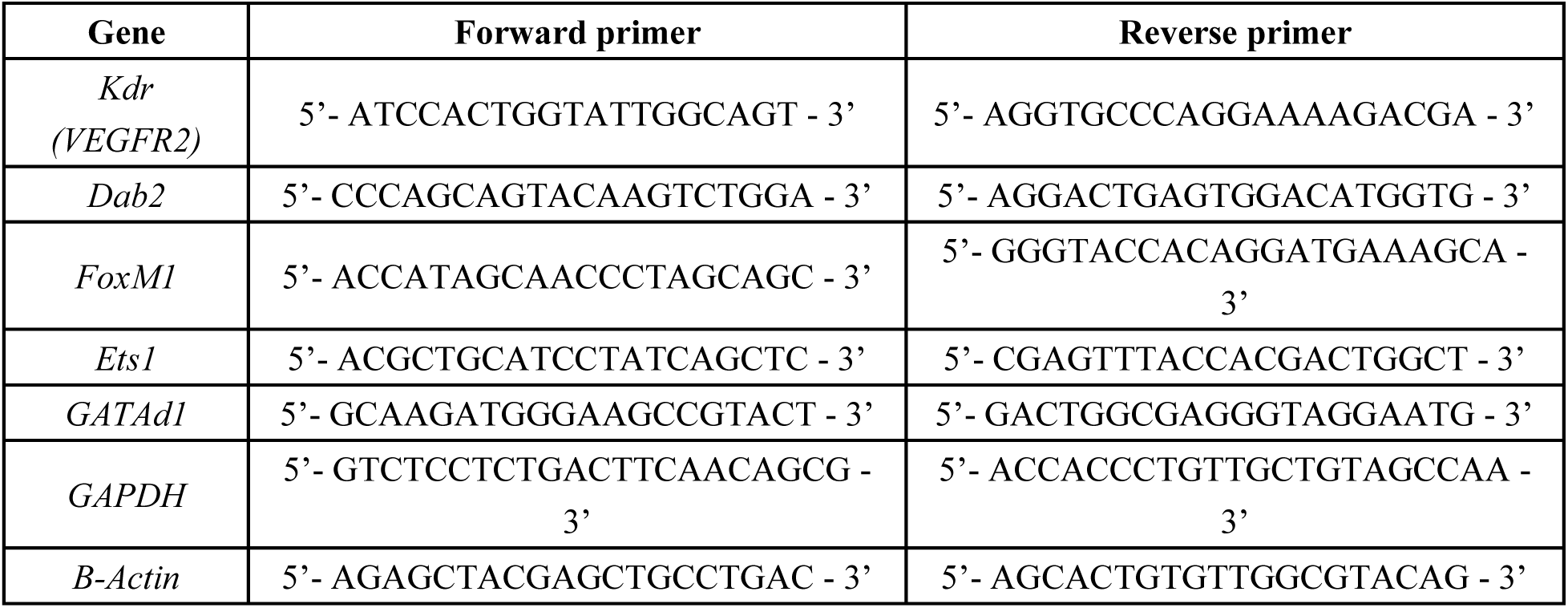
The list of primers used in qRT-PCR.

**Table S2.** The list of antibodies.

| **Target antigen** | **Vendor or Source** | **Catalog #** | **Applications** | **Source Isotype** | **Cross activity** |
| --- | --- | --- | --- | --- | --- |
| VEGFR2 | Cell signaling | 9698 | WB | Rabbit | H M R |
| phospho-VEGFR2 | Cell signaling | 2478 | WB/IF | Rabbit | H M |
| ERK | Cell signaling | 4695 | WB | Rabbit | H M R |
| phospho-ERK | Cell signaling | 9106 | WB | Mouse | H M R |
| Akt | Cell signaling | 9272 | WB | Rabbit | H M R |
| phospho-Akt | Cell signaling | 4058 | WB | Rabbit | H M R |
| Dab2 | Santa Cruz | sc-136964 | WB/IF | Mouse | H M |
| FoxM1 | GeneTex | GTX100276 | WB/IF | Rabbit | H M |
| Actin | Santa Cruz | sc-58673 | WB | Mouse | H M R |
| GAPDH | Santa Cruz | sc-137179 | WB | Mouse | H M R |
| Anti-Mouse IgG (H+L) Secondary Antibody, HRP | Thermo Fisher Scientific | 31430 | WB | Goat | M |
| Anti-Rabbit IgG (H+L) Secondary Antibody, HRP | Thermo Fisher Scientific | 31460 | WB | Goat | R |
| CD31 | Thermo Fisher | BDB550274 | IF | Rat | H M R |
| Alexa Fluor 488 anti-Rat (H+L) | Invitrogen | A-21208 | IF | Donkey | Rat |
| Alexa Fluor 488 anti-Mouse (H+L) | Invitrogen | A-21202 | IF | Donkey | M |
| Alexa Fluor 488 anti-Rabbit (H+L) | Invitrogen | A-21206 | IF | Donkey | R |
| Alexa Fluor 594 anti-Rat (H+L) | Invitrogen | A-21209 | IF | Donkey | Rat |
| Alexa Fluor 594 anti-Mouse (H+L) | Invitrogen | A-21203 | IF | Donkey | M |
| Alexa Fluor 594 anti-Rabbit (H+L) | Invitrogen | A-21207 | IF | Donkey | R |

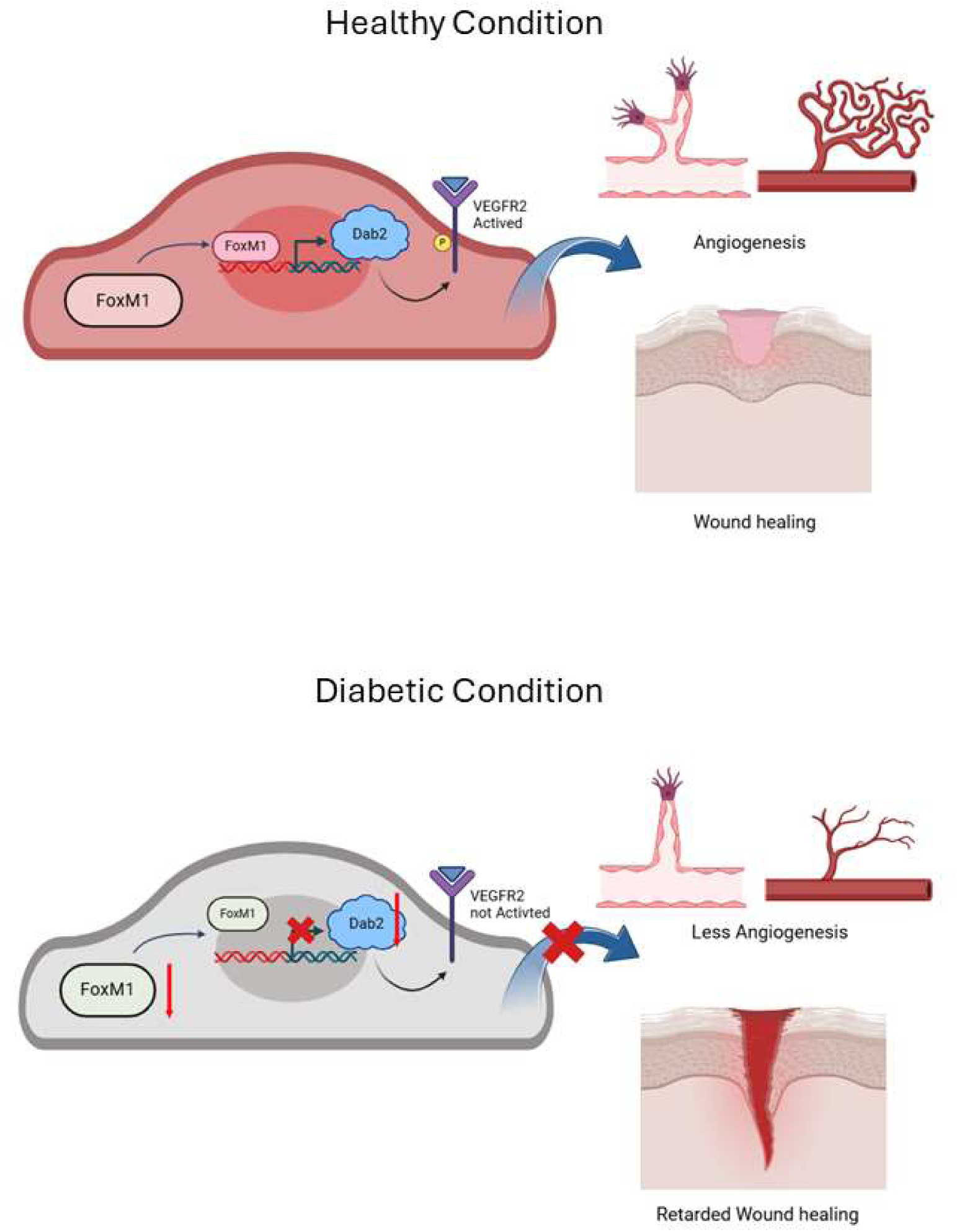

